# Bacterial vitamin sharing emerges from a balance between release and uptake

**DOI:** 10.64898/2026.09.01.748612

**Authors:** Freddy Bunbury, Thomas Janas, Paige Mullen, Kaylie Scorza, Sagnik Ghosh, Jeffrey Zhang, Madhav Mani, Catherine A Pfister, Claire Donnat, Seppe Kuehn

## Abstract

Many microbes use vitamins they cannot synthesize. As a result, vitamin availability impacts microbial community composition and productivity. Yet how vitamins become available to users remains poorly understood. To understand the basis of vitamin sharing, we quantified vitamin B_12_ synthesis and uptake in over 200 soil, freshwater, and marine bacterial isolates. We reveal the existence of four strategies demarcated by all combinations of high or limited B_12_ synthesis and uptake. B_12_ synthesis, but not uptake, was predictable from genomes without pathway knowledge. Among synthesizers, the fraction of vitamin that was extracellular varied over 100-fold. Extracellular B_12_ was quantitatively predicted from a dynamic model capturing only release via death and reuptake by living cells. A subset of synthesizers rapidly reclaim released B_12_, resulting in limited sharing, while retainers exhibit limited release or uptake, and providers release without reuptake. Thus, extracellular B_12_ availability is governed not by synthesis alone, but by the balance between release and uptake.

## 1 Introduction

Microbial communities underpin many processes that sustain ecosystems, from primary production to decomposition (Falkowski et al. 2008; Cavicchioli et al. 2019). The interactions that structure these communities are often mediated through chemical exchanges, as microbes both compete for resources and release metabolites that can be used by others (Douglas 2020; Kost et al. 2023). Vitamins may be especially important in such exchanges because, unlike many nutrients that are incorporated into biomass, they function as enzyme cofactors allowing small concentrations to support large amounts of growth (Seth and Taga 2014; Suazo et al. 2026).

One such vitamin is vitamin B_12_ (cobalamin), a member of the cobamide family, which acts as a cofactor for enzymes involved in central carbon and amino-acid metabolism (Shelton et al. 2019). Cobamide synthesis is restricted to bacteria and archaea, and even among bacteria is highly uneven. Approximately 37% of bacteria are predicted to synthesize cobamides, while 86% can use them (Shelton et al. 2019). This implies widespread acquisition of extracellular cobamides. In marine systems, dissolved B_12_ concentrations are often in the low picomolar range (Bannon et al. 2025; Sañudo-Wilhelmy et al. 2012; Barber-Lluch et al. 2021), and B_12_ addition can increase phytoplankton growth and reshape community composition (Koch et al. 2011; Joglar et al. 2021; Kondo et al. 2024; Rahav and Silverman 2022; Joglar et al. 2020; Wienhausen et al. 2022b; Beauvais et al. 2026). In soils, much of the extracellular B_12_ is matrix-associated, and genomic and culture-based evidence also indicates widespread use and exchange between soil bacteria (Hallberg et al. 2024; Alvarez-Aponte et al. 2024, 2025; Lu et al. 2020). In the human gut, B_12_ status and intake have been associated with microbiome composition and function (Guetterman et al. 2022; Degnan et al. 2014b). Together, these observations identify B_12_ as an ecologically important resource whose production and availability can shape microbial communities.

Classic co-culture studies have shown that bacterial B_12_ producers can support strains requiring B_12_ for growth (auxotrophs) (Haines and Guillard 1974; Croft et al. 2005). These experiments establish that B_12_ synthesized by bacteria can become available, but generally do not identify how it leaves producer cells. Moreover, B_12_-mediated interactions are not uniform: direct sharing in algal–bacterial co-cultures appears strongly species-pair specific (Nef et al. 2022; Bunbury et al. 2022), many marine producers do not release enough vitamin to support algae (Sultana et al. 2023), and soil isolates vary in both B_12_ release and their ability to support auxotrophs (Alvarez-Aponte et al. 2024, 2025). A major unresolved question, therefore, is not whether B_12_ mediates microbial interactions, but what determines how much synthesized B_12_ becomes available to other organisms.

There is evidence that B_12_ can leave bacterial producers through lysis and secretion. The bidirectional cobalamin transporter BacA provides a potential route for export from living cells (Nijland et al. 2024), and bacteriophage-mediated lysis provides a demonstrated route for B_12_ release upon death (Sultana et al. 2025; Wienhausen et al. 2024; da Silva Barreira et al. 2026). However, the relative contributions of release from living and lysed cells and the effect of B_12_ reuptake on extracellular pools remain unresolved (Gregor et al. 2025; Sultana et al. 2023; Alvarez-Aponte et al. 2024).

Here, we combine phenotyping, comparative genomics and quantitative modeling in 274 bacterial isolates across four phyla to determine what controls extracellular B_12_ availability. We find that synthesis, uptake, and release upon cell death can largely explain extracellular B_12_ availability. Controlled cell-death and uptake rate experiments in a representative subset of strains parameterize a dynamic model that quantitatively predicts extracellular B_12_. However, some taxa show higher extracellular B_12_ than expected from cell death alone, potentially suggesting a contribution from living cells. Together, these findings establish synthesis, release, and uptake as separable processes that determine bacterial contributions to extracellular B_12_ pools.

## 2 Results

### Bacterial B_12_ synthesis, uptake, and extracellular accumulation define distinct exchange phenotypes

To ask how bacterial B_12_ sharing phenotypes vary across taxa and environments, we assembled a collection of 274 isolates from soil, freshwater and marine environments spanning four phyla (Fig. S1, Table S1). Of these, 241 were either previously whole genome sequenced or sequenced in this study. Following quality control (≥ 90% completeness, ≤ 5% contamination), 231 genomes were retained for phylogenomic analysis with GTDB-Tk (Fig. S2) (Chaumeil et al. 2020). For each of these strains we assessed B_12_ dependence as the ratio of the area under the OD_600_ growth curve with 10 nM added B_12_ to that without added B_12_. None of the isolates exhibited a B_12_-dependent growth phenotype compared to the *E. coli* Δ*metE* positive control (Fig. S3), suggesting none were B_12_ auxotrophs. This was expected as most isolation media lacked B_12_.

We quantified two B_12_-related traits at early (20-h) and late (44-h) time points in 228 sequenced strains under standardized growth conditions (Table S2). Traits included total B_12_ accumulation, comprising intracellular and extracellular pools, and B_12_ uptake over a 1 h period after addition of 500 pM B_12_. 202 strains yielded measurable traits, of which 39% exhibited B_12_ synthesis (*>*50pM), while 52% took up more than half the added B_12_ (*>*250pM) (Fig. 1A). B_12_ synthesis and uptake both exhibited strong phylogenetic signal (Pagel’s *λ* = 0.99 and 0.98, respectively, *p <* 0.001 for both). Nonetheless, the traits were broadly dispersed, with uptake occurring across all classes, and B_12_ synthesis in every class except Bacteroidia.

**Fig. 1.**
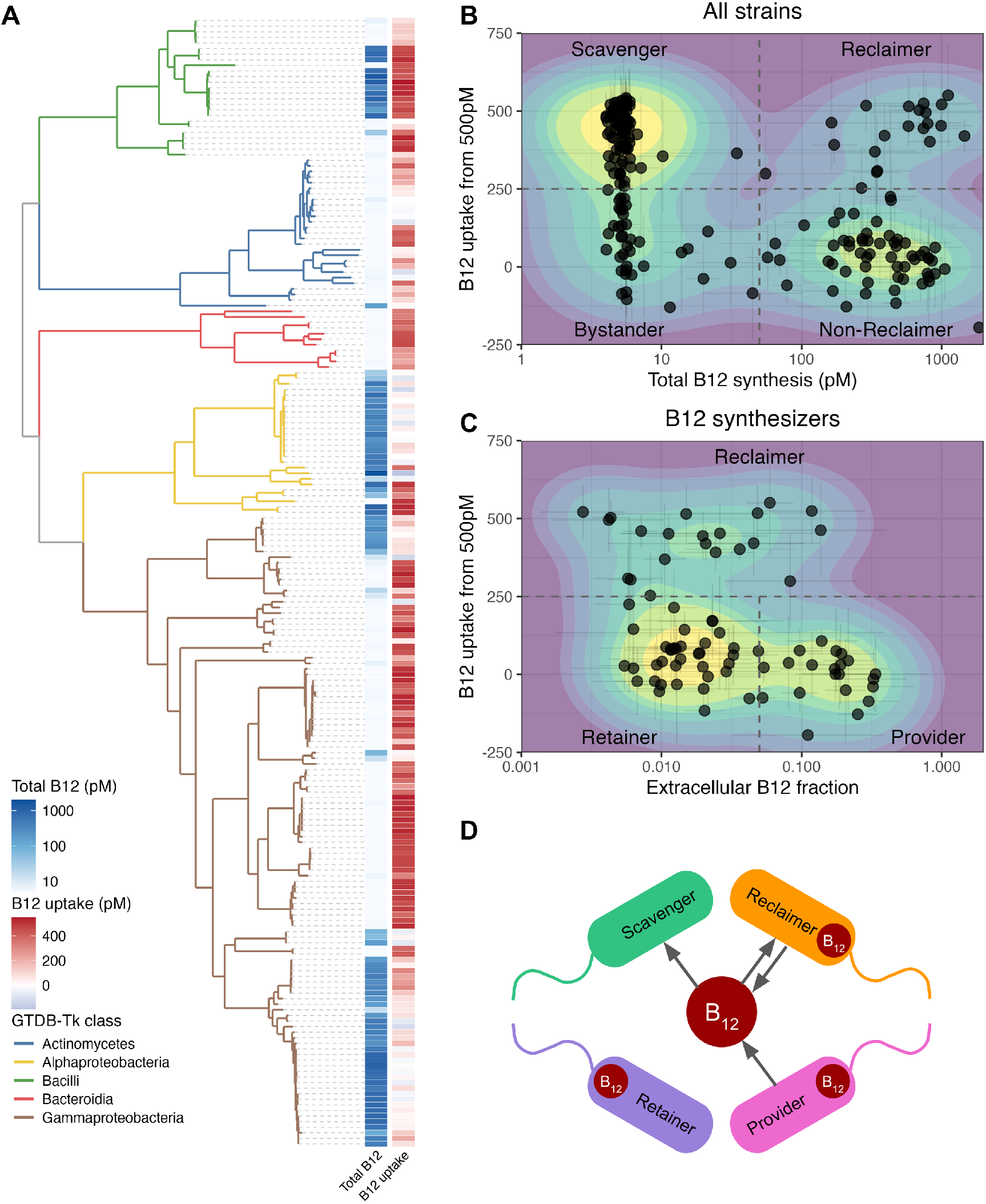
Bacterial B_12_ synthesis, uptake, and extracellular accumulation define distinct exchange phenotypes. **(A)** Phylogenomic tree of the 202 strains with both B_12_ synthesis and uptake measurements, coloured by taxonomic class. Heat map columns show strain-level total B_12_ production and uptake from a 500 pM B_12_ addition on separate colour scales. Values summarize measurements passing the quality criteria using geometric means for total B_12_ and arithmetic means for uptake. **(B)** Relationship between total B_12_ production and uptake across these 202 strains. Each point represents one strain. Dashed lines indicate operational thresholds of 50 pM total B_12_ and 250 pM uptake in one hour, defining bystanders, scavengers, reclaimers and non-reclaimers.**(C)** Relationship between uptake and the fraction of total B_12_ measured extracellularly after 0.2 *µ*m filtration among B_12_ synthesizers (*>* 50 pM total B_12_). Classification was hierarchical: strains with uptake ≥ 250pM were classified as reclaimers irrespective of extracellular fraction. Among non-reclaimers (uptake *<* 250 pM), an extracellular-fraction threshold of 0.05 separated retainers (*<* 0.05) from providers (≥ 0.05). Filled contours in (B) and (C) show bivariate kernel-density estimates. **(D)** Schematic interpretation of the exchange phenotypes. Intracellular B_12_ symbols denote detectable synthesis, inward arrows denote uptake exceeding 250 pM, and outward arrows denote extracellular B_12_ accumulation consistent with externalization. Scavengers take up but do not synthesize detectable B_12_; reclaimers both synthesize and take up B_12_; retainers synthesize B_12_ but show little extracellular accumulation; and providers synthesize B_12_ and accumulate ≥5% extracellularly. Some reclaimers also have extracellular fractions exceeding 0.05 and are therefore depicted with both inward and outward fluxes. Bystanders, which exhibit neither detectable synthesis nor high uptake, and auxotrophs, which were not measured but must take up B_12_, are omitted from the schematic.

Plotting B_12_ synthesis against uptake revealed four trait groups: bystanders (no detectable synthesis and low uptake), scavengers (no detectable synthesis and high uptake), reclaimers (detectable synthesis and high uptake) and non-reclaimers (detectable synthesis and low uptake, Fig. 1B). Scavengers were the most common group (84 strains), followed by non-reclaimers (58), bystanders (39), and finally reclaimers (21). To determine the fraction of synthesized B_12_ that was extracellular, we measured B_12_ after filtering samples through 0.2 *µ*m filters and divided this by the total B_12_ concentration measured in the corresponding unfiltered and lysed sample. Among B_12_ synthesizers, the extracellular B_12_ fraction ranged from 0.3% to 30% of the total (Fig. 1C) and extracellular B_12_ was only weakly positively associated with total B_12_ production (Fig. S4). Among non-reclaimers, which show minimal uptake, extracellular B_12_ fractions formed two clusters. We classified strains with extracellular fractions *<* 5% of the total as retainers and those with fractions ≥ 5% as providers, following previous work on B_12_ sharing (Sultana et al. 2023). Together, these traits, B_12_ synthesis, uptake, and extracellular accumulation, provide an ecological framework for interpreting bacterial contributions to shared B_12_ pools. The likely dominant fluxes into and out of the B_12_ pool associated with each trait group are summarized schematically in Fig. 1D.

### B_12_ synthesis, but not uptake or extracellular accumulation, is predictable from genomes

We inspected the trait groups defined in Fig. 1B,C and found strong association with taxonomic class (Fig. S5; permutation chi-square tests: *p <* 0.001). This motivated us to ask whether B_12_ synthesis, uptake, and extracellular accumulation could be predicted from genomes. We trained random forest classifiers using genome-wide KEGG ortholog presence–absence profiles and compared their classification accuracy with that of a Bernoulli null model based on the prevalence of each trait in the corresponding training set (Fig. 2). Predictions of B_12_ synthesis and uptake were first evaluated across all 202 strains. Since net uptake may be affected by intracellular B_12_ synthesis, we separately evaluated predictions of uptake by synthesizers. Because extracellular accumulation is only relevant for synthesizers we also evaluated predictions for this group.

**Fig. 2.**
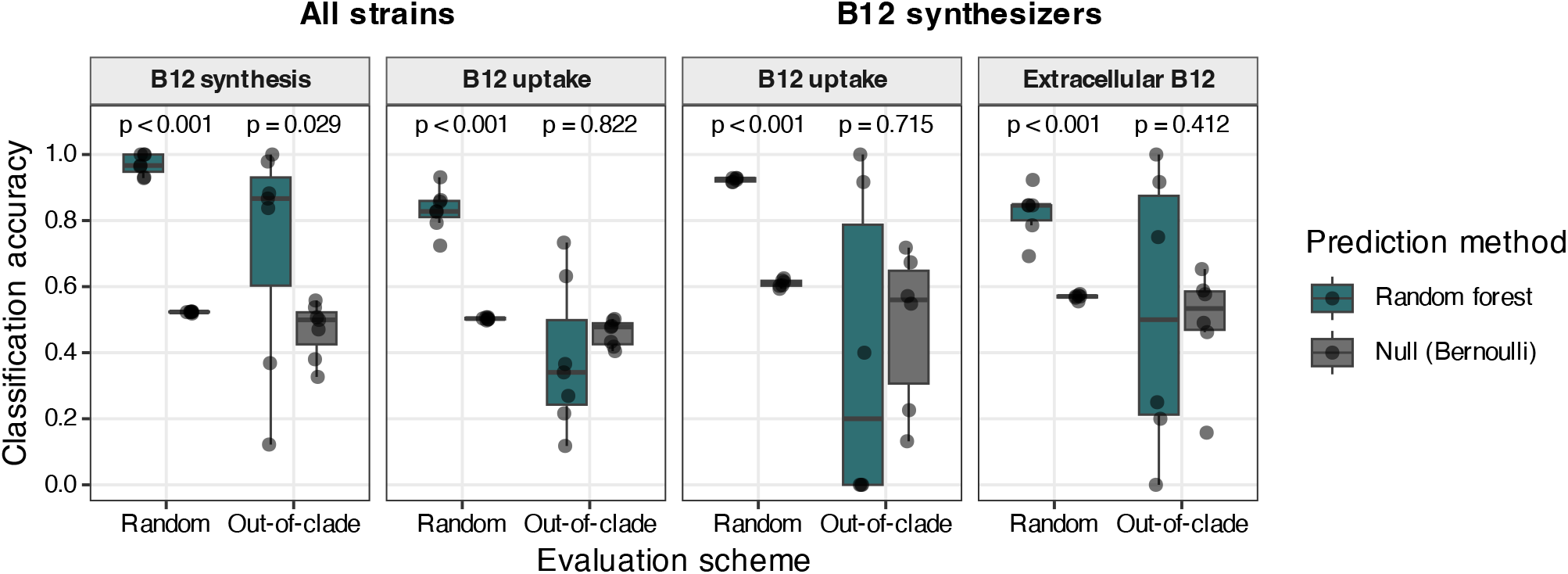
B_12_ synthesis, but not uptake or extracellular accumulation, is predictable from genomes. Random forest classifiers predicted binary B_12_ traits from genome-wide KEGG orthology (KO) presence–absence profiles. Models were evaluated using stratified random test sets (“Random”) or phylogenetically structured test sets formed by holding out a GTDB order, or a block of orders for orders containing *<* 5% of the total genomes (“Out-of-clade”). Models for B_12_ synthesis used all 202 strains, extracellular B_12_ fraction used only B_12_ synthesizers (total *B*_12_ ≥ 50 pM), and B_12_ uptake was assessed using both strain sets. Classification thresholds were total *B*_12_ ≥ 50 pM, B_12_ uptake ≥ 250 pM and extracellular B_12_ fraction ≥ 0.05. Teal boxplots show random-forest classifier accuracy, and grey boxplots show the accuracy of a feature-free null model that randomly assigned classes with probabilities equal to their prevalence in the corresponding training set; points represent individual test splits. Displayed *p*-values are from one-sided paired *t*-tests comparing classifier and null accuracy across matched splits.

When test strains were selected randomly, all traits were predicted significantly more accurately than the null expectation (Fig. 2, Random). However, because closely related strains occurred in both the training and test sets, this predictive signal could reflect either genes directly involved in the traits or phylogenetic correlations in gene content, as discussed by Li et al. (2023).

To distinguish between these possibilities and assess generalization across phylogenetic groups, we repeated the analysis but held out taxonomic GTDB orders as test sets (Fig. S6). Under this more stringent validation scheme, only B_12_ synthesis was predicted significantly better than the null expectation (Fig. 2). Furthermore, several established B_12_ biosynthesis genes were among the most predictive features including genes involved in corrin-ring synthesis and late-stage B_12_ biosynthesis (Fig. S7). We asked whether the genomic features with high importance for random forest predictions (Fig. S7) were also discovered by a phylogenetically controlled association analysis. Using a genome-wide linear mixed model (Lees et al. 2018), we found strong concordance between random-forest feature importance and pyseer association strength for B_12_ synthesis (Fig. S8). By contrast, uptake showed little concordance between random-forest importance and pyseer association strength, and no genes reached significance after multiple-testing correction (Fig. S8). Extracellular B_12_ accumulation similarly lacked clear functionally linked genomic predictors (Fig. S7). Thus, the genomic determinants of B_12_ synthesis appear sufficiently conserved to support prediction across phylogenetic boundaries, whereas uptake and extracellular accumulation may depend more strongly on lineage-specific genes or physiological state. We therefore next asked whether extracellular B_12_ accumulation could instead be explained by physiological processes that release B_12_ into, or remove it from, the extracellular pool.

### B_12_ uptake is correlated with lower B_12_ sharing

Having observed a modest negative association between B_12_ uptake and extracellular accumulation (Fig. S4), we next asked whether cell death could also explain extracellular B_12_ accumulation. We estimated the fraction of dead cells using the dead-cell-specific fluorescent DNA-binding dye DCS1 (AAT Bioquest 50-224-9741). This assay was more sensitive than OD_600_ at detecting low density cultures (Fig. S9A), and highly specific for dead cells, with a linear response to dead cells in live-dead mixtures (Fig. S9B). We found that the dead-cell fraction was not significantly correlated with the fraction of B_12_ found extracellularly (Fig. 3A). Providers combined high extracellular B_12_ fractions with low dead-cell fractions, potentially suggesting an additional non-lytic source of extracellular B_12_. Conversely, many reclaimers fell below the 1:1 expectation if B_12_ released by dead cells remained extracellular, suggesting that released B_12_ was subsequently taken up (Fig. S10). To account for deviations from this 1:1 line we divided the extracellular B_12_ fraction by the dead cell fraction as a proxy for B_12_ sharing. We found that this ratio was negatively correlated with B_12_ uptake (Pearson’s r = -0.577, *p <* 0.001; Fig. 3 B). This association is consistent with uptake reducing the amount of released B_12_ that remains extracellular. Providers therefore appear to be the major sources, whereas reclaimers act as producer-sinks that release B_12_ but also recapture it.

**Fig. 3.**
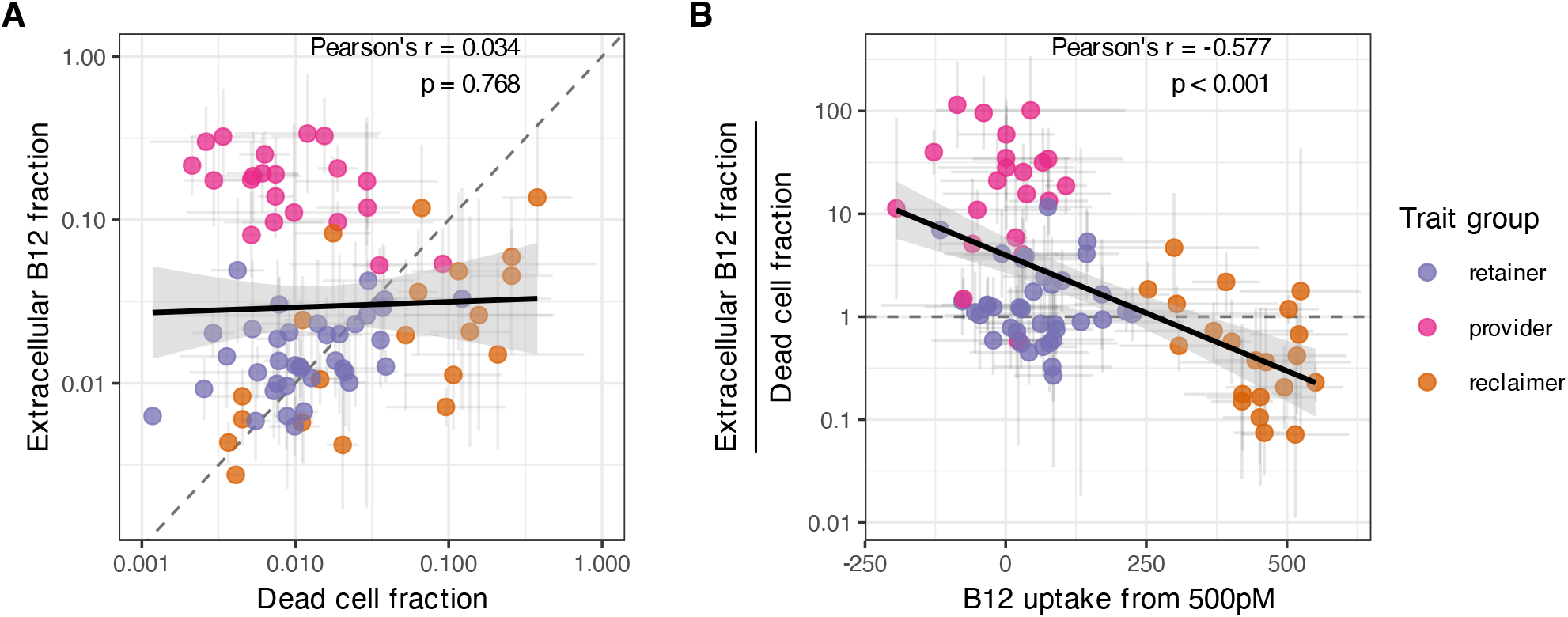
B_12_ uptake is correlated with lower B_12_ sharing. **(A)** Relationship between the extracellular B_12_ fraction (EC-B_12_) and the dead-cell fraction. The dead-cell fraction was estimated from the fluorescence intensity of the dead-cell-specific stain DCS1 applied after sampling, normalized by its intensity after adding it to cultures that were killed by heating at 95 °C for 30 min. The diagonal dashed line indicates the expectation that the EC-B_12_ fraction equals the dead-cell fraction if cell death accounts for all B_12_ release and the released B_12_ remains extracellular. No significant relationship was detected. **(B)** Relationship between B_12_ uptake following a 500 pM addition and the ratio of the EC-B_12_ fraction to the dead-cell fraction. This ratio represents the measured EC-B_12_ fraction relative to that expected from cell death alone; the horizontal dashed line marks a ratio of one. The solid line shows the fitted regression, and the shaded region indicates its 95% confidence interval. The Pearson correlation coefficient (r) and its *p*-value are reported in the upper-right corner. In both panels, points are coloured according to the B_12_ trait groups defined in Fig. 1C. Points show strain means, and error bars show standard deviations, with arithmetic means and standard deviations used on linear scales and geometric versions on logarithmic scales.

### Cell death and uptake quantitatively predict extracellular B_12_

The associations between extracellular B_12_, cell death, and uptake suggested that B_12_ availability might be quantitatively predictable from two processes: release from dead cells and recapture by living cells. To test this, we focused on 23 producer strains across four taxonomic classes, with roughly equal representation of reclaimers, providers and retainers (Table S3).

We first estimated a first-order uptake coefficient, 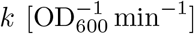, separately for each strain and replicate. The first-order approximation is supported by the relatively low B_12_ concentration used here (500 pM), compared with the apparent *K*_*m*_ of 5 nM reported for the initial phase of B_12_ uptake in *E. coli* (Di Girolamo and Bradbeer 1971). Uptake was modelled as proportional to extracellular B_12_ concentration [pM] and culture density, *x* [OD]:

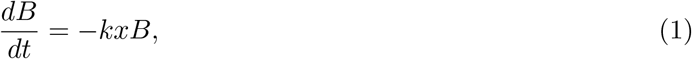

with solution

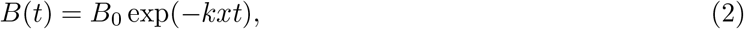

where *B*(*t*) is the blank-adjusted extracellular B_12_ concentration at time *t, B*_0_ is its initial concentration, and *x* is approximated as constant over the four-hour assay using the mean of the initial and final OD_600_, which decreased slightly during the uptake assay (median change of -13%, Fig. S11). Short-term extracellular B_12_ dynamics following the addition of 500 pM B_12_ showed rapid depletion by reclaimers, whereas providers and retainers showed little change (Fig. 4A & S12). Fitting Equation 2 to these dynamics yielded values of *k* that were highest in reclaimers and generally lowest in providers (Fig. 4B). Because B_12_ levels appeared to saturate in some reclaimers, we also tested a two-parameter quota-based uptake model, which resulted in similar findings (Fig. S12).

**Fig. 4.**
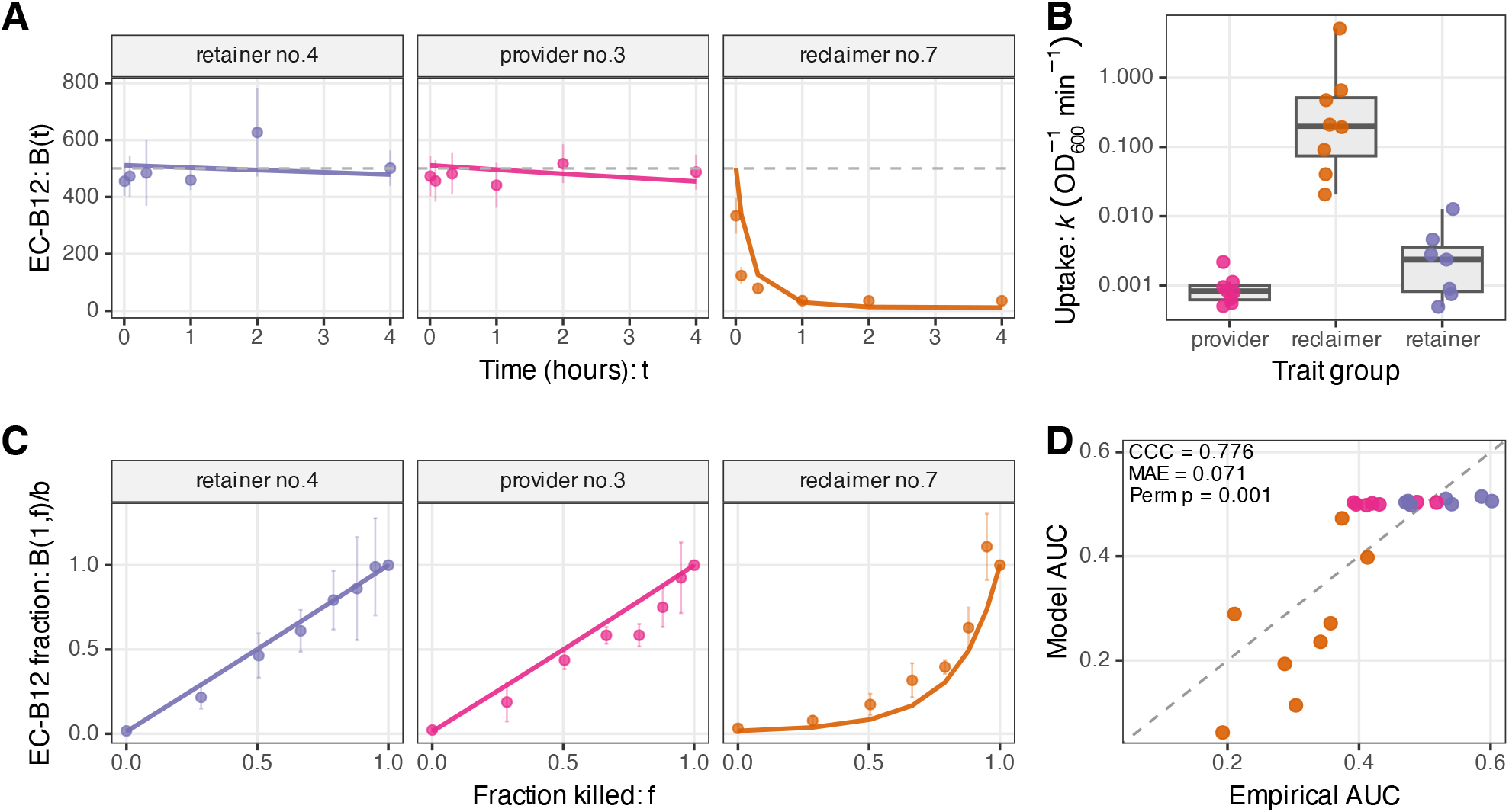
Cell death and uptake quantitatively predict extracellular B_12_. **(A)** Extracellular B_12_ concentrations over 4 h following the addition of 500 pM B_12_ to washed, stationary-phase bacterial monocultures. One representative strain from each B_12_ trait group is shown. Points and error bars represent the mean *±* SD of four replicates. Solid lines show extracellular B_12_ concentrations fitted using Equation 2, and the dashed horizontal line marks the added B_12_ concentration. **(B)** Uptake-rate coefficients (*k*) estimated for all 23 strains. Points represent individual strains, and boxplots summarize the distribution within each trait group. **(C)** Extracellular B_12_ measured 1 h after mixing live and killed cells, expressed as a fraction of the extracellular B_12_ concentration in the corresponding fully killed culture and plotted against the fraction of cells killed. One representative strain from each trait group is shown. Points and error bars represent the mean *±* SD of four replicates. Solid lines show predictions from Equation 3 using independently estimated strain-specific values of *k*. **(D)** Agreement between predicted and empirical responses across all 23 strains. For each strain, the predicted and empirical extracellular B_12_ response curves were summarized as the area under the curve (AUC) of extracellular B_12_ fraction versus fraction killed. The dashed line denotes perfect agreement. Agreement was quantified using Lin’s concordance correlation coefficient (CCC) and the mean absolute error (MAE). A strain-label permutation test assessed whether the observed strain-specific predictions produced a lower MAE than random assignments of model AUCs among strains. Points and lines are coloured consistently by B_12_ trait group across panels.

We then asked whether these independently estimated uptake rates could predict extracellular B_12_ availability following cell death. We assumed that dead cells instantaneously release their intracellular B_12_, establishing an initial extracellular concentration proportional to the dead-cell fraction, while surviving cells recapture extracellular B_12_ according to the uptake coefficient *k* and surviving culture density *x*(1 − *f*). This gives:

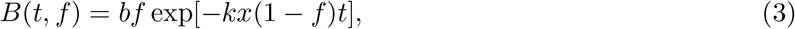

where *f* is the dead-cell fraction and *b* is the B_12_ concentration in the fully heat-killed culture. To control *f* experimentally, we split stationary-phase cultures into two aliquots, heat-killed one aliquot and remixed live and killed cells in defined proportions. After 1 h, cultures were filtered and extracellular B_12_ was measured.

In this controlled-death assay, equation 3 correctly predicted extracellular B_12_ after one hour of incubation. Using the independently estimated strain-specific values of *k*, we predicted *B*(1*h, f*) ≈ *bf* for providers and retainers consistent with their near zero uptake coefficients *k* (Fig. 4C & S13). Reclaimers, however, left less B_12_ extracellularly, consistent with rapid recapture by surviving cells (Fig. 4C & S13). We found similar results for the quota-based model (Fig. S13).

To assess the quality of these predictions we computed the area under the curve (AUC) describing extracellular B_12_ fraction as a function of fraction of cells killed. In the absence of uptake, the expected AUC is ~0.5, whereas increasing uptake lowers it toward 0. We compared AUC for the data to predictions from the model. The model predicted AUC values close to 0.5 for providers and retainers and lower values for reclaimers, as we observed (Fig. 4 D). Predicted and empirical AUCs showed strong concordance (CCC = 0.776), and a low mean absolute error (MAE = 0.071) (Fig. 4 D). Moreover, the observed strain-specific predictions had a significantly lower MAE than randomly reassigning model predictions among strains (1,000-permutation test, *p* = 0.001; Fig. 4 D). We found similar results when using the quota model to describe uptake (Fig. S13). Thus, extracellular B_12_ availability can be quantitatively predicted from a balance between release through cell death and recapture by living cells.

### The balance between release and uptake predicts extracellular B_12_

Release of B_12_ by dead cells and re-uptake by living cells explained much of the extracellular levels observed in the live–dead mixtures (Fig. 4D). However, in earlier measurements, providers accumulated more extracellular B_12_ than expected from their measured dead-cell fractions suggesting a non-lytic route for B_12_ release (Fig. 3A). We therefore asked whether stimulating growth through glucose addition might increase the amount of extracellular B_12_. We measured short-term extracellular B_12_ dynamics after washed cultures were resuspended in B_12_-free medium with or without 1.67 mM added glucose. Providers showed the greatest increases in extracellular B_12_ over time, particularly in the presence of glucose, whereas retainers and reclaimers showed comparatively less accumulation (Fig. 5A & S14).

**Fig. 5.**
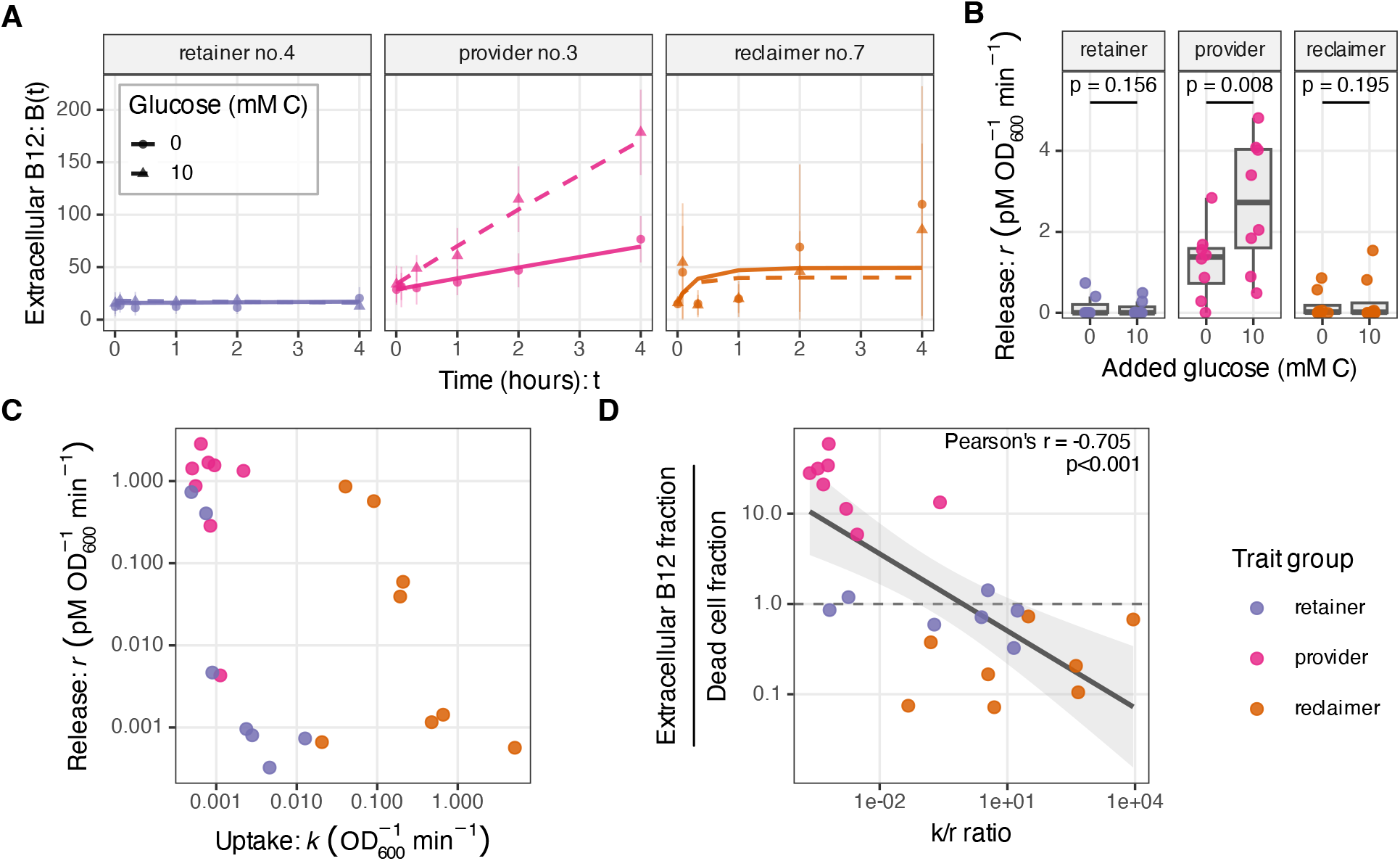
The balance between release and uptake predicts extracellular B_12_. **(A)** Extracellular B_12_ concentrations over 4 h after washed bacterial cultures were resuspended in medium with or without 1.67 mM added glucose (10 mM of carbon atoms). Lines show extracellular B_12_ concentrations fitted using Equation 5. For each strain and glucose condition, the uptake-rate coefficient (*k*) was fixed at its independently estimated value from Fig. 4, while the release-rate coefficient (*r*) was fitted to the data. Fixing *k* at its independently estimated value reduced trade-offs between the release and uptake coefficients. **(B)** Release-rate coefficients (*r*) estimated for all 23 strains in the absence vs presence of glucose. Annotated *P* -values are from two-sided exact paired Wilcoxon signed-rank tests comparing the two glucose treatments within each trait group. **(C)** Relationship between the release-rate (*r*) and uptake-rate (*k*) coefficients across the 23 strains. Each point represents one strain, with *r* and *k* calculated as geometric means of replicate-specific OD-normalized coefficients in the absence of glucose. **(D)** Extracellular B_12_ availability relative to cell death, expressed as the extracellular B_12_ fraction divided by the dead-cell fraction, plotted against the ratio of uptake to release (*k/r*). Each point represents one of the 23 strains for which both *k* and *r* were estimated; extracellular B_12_ and dead-cell fractions are from Fig. 3A. The dashed horizontal line at *y* = 1 denotes the null expectation that the extracellular B_12_ fraction equals the dead-cell fraction. The solid line shows a linear regression fitted to the log_10_-transformed ratios, and the shaded region indicates its 95% confidence interval. Pearson’s correlation coefficient and its associated *p*-value are shown in the upper-right corner. Points and lines are coloured consistently by B_12_ trait group across panels.

To quantify B_12_ release while accounting for its simultaneous uptake, we added a constant release-rate coefficient, *r*, to the uptake model:

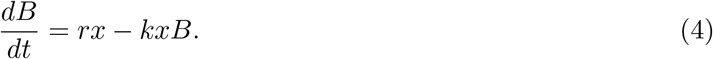

Solving Equation 4 with the initial condition *B*(0) = *B*_0_ gives

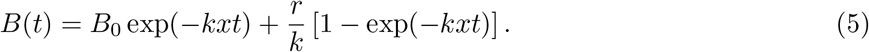

Here, *B*(*t*) is the extracellular B_12_ concentration at time *t, B*_0_ is its initial concentration, *r* is the release-rate coefficient, and *k* is the uptake-rate coefficient. We assumed release was constant per unit OD_600_ because intracellular B_12_ concentrations can greatly exceed extracellular concentrations (Kennedy et al. 2022; Sultana et al. 2023) and producers commonly regulate B_12_ biosynthesis through riboswitches (Moore et al. 2014; Vitreschak et al. 2003), such that this concentration difference is expected to change little over the assay. We use “release” and “uptake” as operational descriptions of the positive and negative fluxes represented by the model; other processes that increase or decrease extracellular B_12_ may also contribute to these coefficients.

For each strain and glucose condition, we fitted Equation 4 while fixing *k* at the value estimated from the uptake experiment in Fig. 4. Fixing *k* at its independently estimated value reduced trade-offs between the release and uptake coefficients. This approach aids in identifying both rates because we manipulate the extracellular B_12_ levels, which are high in the uptake experiment (*kxB >> rx*) and low in the release experiment (*kxB << rx*). However, a B_12_ labeling experiment would be required to make a definitive statement on absolute rates of uptake and release.

Providers were the only group showing a significant increase in the fitted release-rate coefficient following glucose addition (Fig. 5B). However, on average glucose addition increased culture density and dead-cell signal in providers (Fig. S15). Extracellular B_12_ normalized by dead-cell signal did not change significantly in providers, but decreased significantly in retainers and reclaimers (Fig. S15). Thus, growth by retainers and reclaimers stimulates net B_12_ uptake. For providers, while we observe net release on stimulation with glucose, we cannot say definitively that this release arose from living cells.

Combining the fitted release and uptake-rate coefficients separated the three trait groups in release/uptake space: providers generally combined high release with low uptake, retainers combined low release with low uptake, and reclaimers combined high uptake with variable release (Fig. 5C, S16).

We next asked whether the balance between uptake and release predicted extracellular B_12_ availability relative to measured cell death using the data presented in Fig. 3. Across the 23 strains, the ratio of the extracellular B_12_ fraction to the dead-cell fraction decreased significantly as the uptake-to-release ratio (*k/r*) increased (Fig. 5D). Thus, strains with high release relative to uptake provided more extra-cellular B_12_ than expected from cell death, whereas strains with high uptake relative to release left less B_12_ available.

## 3 Discussion

We aimed to determine why B_12_-producing bacteria differ in the amount of vitamin they share. Across a broad collection of environmental isolates, we found that B_12_ synthesis alone was a poor proxy for extracellular availability (Fig. 1C). By combining B_12_ synthesis, uptake, and extracellular accumulation traits, we defined source–sink strategies: scavengers acquired B_12_ without producing it and therefore acted as strong sinks; providers left B_12_ available extracellularly and therefore acted as strong sources; and reclaimers both released and recaptured B_12_, making their net contribution dependent on the balance of these processes. While prior studies and reviews have defined B_12_ functional groups and observed that producers vary in the amount of B_12_ they share (Droop 2007; Sultana et al. 2023; Alvarez-Aponte et al. 2025; Bertrand et al. 2026), our study shows that understanding B_12_ sharing requires dynamic characterization of release and uptake.

Whether B_12_ leaves microbial cells through lysis or release from living cells has long been discussed (Carlucci and Bowes 1970; Haines 1974; Droop 2007). Viral-mediated lysis increases B_12_ release in laboratory culture (Wienhausen et al. 2024; Sultana et al. 2025; da Silva Barreira et al. 2026), and is likely a major route for B_12_ sharing in the environment (Bertrand et al. 2007; Wienhausen et al. 2022a). In our experiments, the dead cell and extracellular B_12_ fractions varied considerably but were low on average (geometric mean), at 1.5% and 3%, respectively (Fig. 3A). Additional stressors could therefore increase extracellular B_12_ pools substantially. However, our findings add a quantitative refinement: cell death may contribute to the B_12_ pool, but reuptake counteracts this.

On the other hand, providers showed more extracellular B_12_ than expected from measured dead cells alone. They also showed increases in B_12_ release during growth (Fig. 5B); however, these were also correlated with increased dead cell abundance (Fig. S15). We therefore cannot definitively say whether our results support other observations that B_12_ provision can occur while producers are alive (Kazamia et al. 2012; Grant et al. 2014; Iguchi et al. 2011; Malalasekara and Escalante-Semerena 2026). A candidate route for B_12_ export is BacA, a multi-substrate transporter shown to transport B_12_ bidirectionally (Nijland et al. 2024). In our data, BacA presence was associated with lower net B_12_ uptake (Fig. S17); however, this was not significant after correcting for population structure, so a causal role requires direct testing.

Modeling approaches have found that vitamin auxotrophs can persist with producers under both lytic and secretion scenarios (Gregor et al. 2025). Importantly, however, our results suggest that B_12_-independent species, such as scavengers and reclaimers, may compete with auxotrophs for access to the extracellular B_12_ pool.

We found that genome-based predictions of B_12_ synthesis are accurate across a broad set of bacteria, while predicting sharing remains limited (Fig. 2). Our statistical approaches identified corrin ring and lower ligand synthesis (*bluB*) genes as the best predictors of measured B_12_ synthesis (Fig. S8). By contrast, B_12_ uptake was harder to predict (Fig. S7). Transporter annotations remain a weak point in predicting microbial metabolic interactions, because errors in substrate assignment, complex membership, or directionality can affect inferred exchange (Casey et al. 2024; Degnan et al. 2014a). Our results also showed that uptake varied with glucose addition (Fig. S15), indicating that it depends on physiological context. Thus, producer reuptake, one of the most important traits for realized sharing, appears variable and remains difficult to infer from current genome annotations.

Across environments, extracellular B_12_ can become less accessible after it is released. In marine systems, B_12_ may be diluted or rapidly degraded by light (Bannon et al. 2024; Bertrand et al. 2007; Sañudo-Wilhelmy et al. 2012; Bannon et al. 2025); in grassland soil, extracellular corrinoids are largely matrix-associated (Hallberg et al. 2024); and in the gut, B_12_ can be bound by extracellular vesicles (Juodeikis et al. 2022, 2025). Nevertheless, microbial use of exogenous cobamides is widespread (Croft et al. 2005; Shelton et al. 2019; Lu et al. 2020). Local exchange within communities may allow microbes to intercept released vitamins before they are diluted, degraded, or otherwise made inaccessible. A community’s capacity to support B_12_-dependent taxa may therefore depend not on biosynthetic potential alone, but on the physical structure of its environment and the balance between microbial release and uptake.

### Limitations of this study

We quantified B_12_ using an algal bioassay rather than chemically resolving individual cobamides by liquid chromatography–mass spectrometry (Mok et al. 2022). This functional measure suits our focus on the biological consequences of sharing but reflects availability to the assay organism, not total cobamide abundance.

Our dynamic extracellular B_12_ measurements cannot fully resolve simultaneous release and uptake, particularly when these fluxes are similar in magnitude. However, we partially separated these processes by measuring dynamics with and without added B_12_, under a first-order uptake assumption.

We have referred to extracellular B_12_ accumulation in bacterial monocultures as a proxy for sharing without directly demonstrating uptake by non-producers in coculture. Future experiments combining producers and auxotrophs in cocultures could test whether measured extracellular availability predicts realised B_12_ transfer and support of auxotroph growth.

Finally, our study does not experimentally establish the molecular mechanisms underlying B_12_ release and uptake. Future work could test whether candidate transporters promote or limit B_12_ sharing across these strains using gene knockouts and complementation, as has been done in model B_12_ producers such as *S. typhimurium* (Malalasekara and Escalante-Semerena 2026).

## 4 Methods

### 4.1 Bacterial isolate collection and cultivation

The experiments used 274 unique bacterial isolates from soil, freshwater, and marine environments, drawn from the collection documented in Table S1. Of these, 67 soil isolates were obtained as part of this study; the remaining 207 were used in previously published studies or had genomes previously deposited at NCBI. Briefly, soil samples were suspended by vortexing in sterile 0.9% w/v NaCl, serially diluted into 10% tryptic soy broth (TSB) in 96-well plates, and incubated at 25 °C for 48 h. Selected cultures were streaked onto 10% TSB 1.5% agar, and individual colonies were picked to establish isolate cultures. The full set of isolates was arrayed into three 96-well plates for downstream handling.

Soil and freshwater isolates in arrays 1 and 2 were cultured in 10% TSB for 48 h, whereas marine isolates in array 3 were cultured in Marine Broth M13 for 96 h; cultures were grown in deep-well 96-well plates. Media recipes are given in Table S2. Glycerol stocks were prepared by combining 25 *µ*L of culture with 25 *µ*L of 40% w/v glycerol in round-bottom 96-well polystyrene plates. Plates were sealed with adhesive PCR plate foil (Thermo Scientific, AB-0626) and stored at −80 °C.

For revival, 100 *µ*L of TM-mod medium was added to each 50 *µ*L glycerol stock to thaw and resuspend the culture. A 10 *µ*L aliquot was then transferred into 990 *µ*L of TM-mod medium supplemented with soytone (25 mg L^−1^), glucose (1.67 mM), vitamin solution A (no B_12_) (0.1%v/v) and methionine (10 *µ*M). Precultures were incubated in the dark for 72 h at 25 °C with shaking at 500 rpm on a microplate shaker (Fisherbrand, 02-217-759) before use in experiments.

### 4.2 B_12_ requirement testing

Seventy-two-hour precultures, prepared as described above, were inoculated in technical duplicate at a 250-fold dilution into flat-bottom 96-well culture plates containing a final volume of 200 *µ*L per well. The basal medium was Tm-mod supplemented with soytone (25 mg L^−1^) and glucose (1.67 mM). Four medium formulations comprised all combinations of methionine (0 or 100 *µ*M) and cyanocobalamin (0 or 10 nM). Plates were incubated in a Liconic Instruments STX44-HRBT automated bench-top incubator operating at 25° C, 95% relative humidity and 800rpm for approximately 57 hours. Plates were individually transferred between the incubator and a BMG CLARIOstar for OD_600_ measurements approximately every 93 minutes by a Hudson PlateCrane EX robotic arm.

B_12_-dependent growth was assessed using cultures grown without added methionine. For each strain, the response to B_12_ was calculated as the ratio of the mean area under the OD_600_-versus-time curve (AUC) with 10 nM cyanocobalamin to the mean AUC without added cyanocobalamin. Because only two replicate wells were available per treatment, comparisons were descriptive and no strain-level hypothesis tests were performed. Responses were compared with that of the B_12_-dependent positive control, *Escherichia coli* Δ*metE*.

### 4.3 Experimental setup for survey of bacterial total B_12_, extracellular B_12_, B_12_ uptake, growth, and cell death

Seventy-two-hour precultures were inoculated at a 500-fold dilution into 1 mL of TM-mod medium supplemented with soytone (25 mg/L) and glucose (1.67 mM) in 96-well deep-well plates. These were sealed with breathable plate membranes (Breathe-Easy, Sigma Z763624) and incubated at 25 °C with shaking at 500 rpm in the dark. Samples of 300 *µ*L were taken aseptically after 20 and 44 h of culturing. These samples were aliquoted for measurements of total B_12_, extracellular B_12_, B_12_ uptake, OD_600_, and dead-cell abundance.

For B_12_-uptake measurements, B_12_ was added at a final concentration of 500 pM to one aliquot in a round-bottom 96-well plate, which was incubated as above for 1 h. The original unamended culture and the 500-pM-amended culture after 1 h were vacuum-filtered through 0.2-*µ*m 96-well filter plates (Fisher Scientific MSGVN2210) using a vacuum manifold, with the filtrate collected in 96-well polystyrene plates. One aliquot of the unamended filtrate was then spiked to 500 pM B_12_ to serve as the “0-h” uptake sample. This prevented uptake during the unavoidable first 30 s after spiking and before filtration from reducing the measured extracellular B_12_. The three filtrate plates and one total-B_12_ plate were then sealed with adhesive PCR plate foils (Thermo Scientific AB-0626) and incubated at 95 °C in an oven for 30 min to lyse bacterial cells in the total-culture aliquot and sterilize all fractions. Samples were then stored at −80 °C for the B_12_ bioassay described below.

OD_600_ was measured in technical duplicate using 45-*µ*L culture aliquots in 384-well plates. The second replicate plate was then sealed as above, incubated at 95 °C for 30 min to kill the cells, and cooled to room temperature. To quantify dead cells, we used the dead-cell-specific fluorescent dye Nuclear Green DCS1 (AAT Bioquest 50-224-9741). This was added to the untreated and heat-treated aliquots at a final 2,000-fold dilution (2.5 *µ*M), and the plates were incubated in the dark. Fluorescence was measured after 15 and 20 min using a VANTAstar plate reader (BMG LABTECH), with excitation centered at 503 nm with a 10-nm bandwidth and emission centered at 538 nm with a 10-nm bandwidth. The two measurement time points confirmed minimal change in signal and were averaged. The fraction of cells that were dead in each culture was calculated by dividing the fluorescence of the untreated sample by that of the heat-treated sample. Two experimental replicates were performed.

### 4.4 B_12_ quantification using an algal bioassay

The cobalamin bioassay used the B_12_-dependent *Chlamydomonas metE4* strain generated by Bunbury et al. (2020) and was similar to the assay described by Harrison et al. (2026). The assay was validated by comparison against a cobalamin bioassay based on *Escherichia coli* K-12 MG1655 (NRRL B-65639; ATCC 700926) and the isogenic derivatives Δ*metE* (NRRL B-65640) and Δ*metE* Δ*metH* (NRRL B-65641), as described by Mok et al. (2022), but modified for 384-well plates (Fig. S18).

Three days before each assay, the *metE4* inoculum was prepared by diluting a maintained late-log-phase culture 100-fold into 1 mL of TAP medium supplemented with 200 pM B_12_. The inoculum was cultured at 28 °C under 80 *µ*mol photons m^−2^ s^−1^ white light without shaking. Before inoculation, this culture was diluted in B_12_-free TAP assay medium to give a final 2,000-fold dilution of the inoculum in each assay well. Each B_12_ sample was added at both 10 and 30 *µ*L to assay medium predispensed into a 384-well plate, giving a final volume of 100 *µ*L per well. Standards spanning 0–5,000 pM before addition to the assay were similarly added at both 10 and 30 *µ*L, producing final in-well B_12_ concentrations ranging from 0 to 1,500 pM (lowest nonzero concentration, 0.0524 pM). Standards were prepared in either blank TM-mod medium or 40-h culture filtrate from three phylogenetically diverse bacterial non-producers of B_12_, partially accounting for nonspecific effects of the sample matrix on the bioassay; these effects were minimal. The plates were lidded and incubated upside down at 28 °C under 80 *µ*mol photons m^−2^ s^−1^ white light without shaking. Chlorophyll fluorescence was measured on days 3–4 using a VANTAstar plate reader (BMG LABTECH), with excitation centered at 400 nm with a 100-nm band-width and emission centered at 700 nm with a 100-nm bandwidth. Standard curves and distributions of sample values are shown in Fig. S19.

Raw fluorescence values were log transformed and converted to B_12_ concentrations using a custom R pipeline. Significant systematic edge effects in the outer rows of the bioassay plates were corrected using paired standards from adjacent inner rows. Four-parameter logistic curves relating log fluorescence to log B_12_ concentration were fitted separately for each assay batch, measurement time, and bacterial source used to prepare the standards:

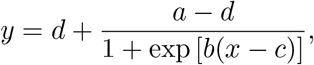

where *y* is log fluorescence, *x* is log B_12_ concentration, *a* and *d* are the upper and lower response asymptotes, respectively, *c* is the log concentration at the midpoint, and *b* describes the slope of the curve. Sample concentrations were estimated jointly from all applicable curves and assay dilutions, with curves weighted according to their residual variance. Estimates were corrected for assay and storage dilution and reported in picomolar units relative to the original sample. Estimation was restricted to the concentration range represented by the contributing standard curves, and fluorescence measurements outside the central 90% of a fitted curve’s response range were bounded at the corresponding limit. One-hour B_12_ uptake was calculated by subtracting the 1-h concentration from the baseline concentration. The baseline was defined as the mean of the measured 0-h concentration following B_12_ addition and the expected concentration, calculated as the corresponding no-addition measurement plus 500 pM. Where indicated, B_12_ measurements were normalized by blank-corrected OD_600_. Samples were retained for downstream trait analyses when their OD_600_ exceeded the blank mean by two standard deviations.

### 4.5 Short-term extracellular B_12_ release and uptake dynamics

Twenty-three selected B_12_-producing strains were cultured and stored as two biological-replicate glycerol stocks (Table S3). These strains were cultured in 1 mL of TM-mod medium + soytone (25 mg/L) + glucose (1.67 mM) for 72 h as before, and then inoculated at a 100-fold dilution into 8 mL of medium in 15 mL centrifuge tubes (Falcon 05-527-90) and incubated at 25 °C and 200 rpm for 20 h. The tubes were centrifuged at 3500 × *g* for 5 min, vortexed for 5 s to loosen the pellet, and resuspended in 1 mL TM-mod. For the B_12_ uptake assay, this eightfold-concentrated suspension was diluted eightfold by addition to TM-mod medium + soytone (25 mg/L) + 500 pM B_12_, either with or without glucose (1.67 mM), pipetted up and down three times, and then immediately vacuum-filtered through a 0.2-*µ*m filter plate as before. The assay plate was then incubated at 25 °C with shaking at 500 rpm. Aliquot filtering was repeated at 5, 20, 60, 120, and 240 min after mixing. The B_12_ release assay was performed similarly, except that strains were added to medium lacking B_12_. Filtered aliquots were sealed, heated at 95 °C for 30 min, and then stored at −80 °C as before. OD_600_ and dead-cell quantification using DCS1 were performed as before at 0 and 240 min. Two experimental replicates, each with two biological replicates, were performed.

### 4.6 Controlled live–dead cell mixing experiment

The live–dead mixing experiment used the same eightfold-concentrated resuspensions of 23 B_12_-producer strains as the release and uptake dynamics experiment described above. One aliquot was sealed in a 96-well plate and heated at 95 °C for 30 min to kill the cells, while the other was left untreated. Both were then diluted eightfold in TM-mod to restore the starting culture concentration. The live and dead fractions were mixed in different ratios to produce cultures containing 0%, 28.5%, 50.5%, 66.5%, 79%, 88%, 95%, or 100% dead cells. These cultures were incubated at 25 °C with shaking at 500 rpm for 1 h, then vacuum-filtered through 0.2-*µ*m filters. Unfiltered aliquots were used for dead-cell quantification, and filtered aliquots were heated and stored for the B_12_ bioassay.

### 4.7 Modelling extracellular vitamin B_12_ uptake and release

Extracellular vitamin B_12_ dynamics were modelled in R using a two-stage kinetic framework. Culture-specific models were fitted separately for each bacterial strain, glucose treatment, experiment, and biological replicate. Because the same cultures were used in the release-and-uptake and live–dead mixture experiments, parameters estimated from the release-and-uptake time courses were used to predict extracellular B_12_ concentrations in the corresponding live–dead mixtures. We express the models using culture density *x*, represented by blank-adjusted OD_600_ and treated as constant during each time course, and OD-specific uptake and release coefficients *k* and *r*. For the main first-order analysis, uptake and release coefficients were normalized to the mean of the blank-adjusted OD_600_ measured at 0 and 240 minutes for each biological replicate and glucose condition. Initial OD was measured in the untreated source culture; final OD was averaged across the corresponding cultures with and without added B_12_. Fig. 4B, 5B&C, and S16 show geometric means of these OD-specific coefficients across replicates. First-order models were fitted to log-transformed B_12_ concentrations. B_12_ concentrations from the bioassay measurements were likewise estimated on a logarithmic concentration scale (see above).

A single blank offset, *B*_blank_, was calculated as the mean B_12_ concentration across all blank-culture samples receiving no added B_12_. The uptake-rate coefficient was estimated from time courses receiving 500 pM added B_12_, using

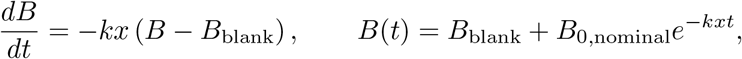

where *B*(*t*) is the predicted extracellular B_12_ concentration at time *t*, and *B*_0,nominal_ is the concentration of added B_12_ (500 pM). Here *k* has units of 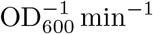; the fitted culture-level coefficient was constrained to 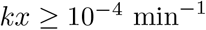.

The release-rate coefficient was subsequently estimated from the corresponding time course without added B_12_. Because release and uptake occurred concurrently, *k* was fixed at the value estimated from the added-B_12_ time course, allowing release to be estimated independently of uptake. The initial extracellular concentration was fixed from the earliest measurement (15 s):

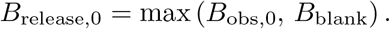

Release was assumed to occur at a constant rate, and extracellular B_12_ was modelled as

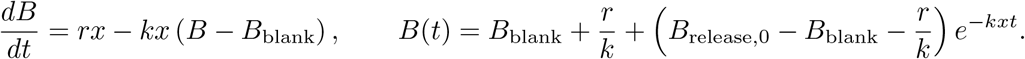

Here *r* has units of 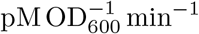; the fitted culture-level release coefficient was constrained to *rx* ≥ 10^−4^ pM min^−1^.

#### 4.7.1 Prediction of extracellular B_12_ in live–dead mixtures

The amount of extracellular B_12_ associated with complete cell killing, *b*, was estimated for each strain, glucose treatment, experiment, and biological replicate from the fully killed bacterial cultures. For a live–dead mixture with killed-cell fraction *f*_dead_, the corresponding live-cell fraction was defined as *f*_live_ = 1 − *f*_dead_, and the initial extracellular B_12_ concentration was predicted as

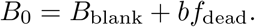

Extracellular B_12_ remaining after uptake by the live fraction of the culture was predicted at *t* = 60 min using the uptake coefficient estimated from the corresponding added-B_12_ time course. The release-rate coefficient was not included in these main-figure predictions; thus, the model attributed changes after mixing solely to uptake by the remaining live cells:

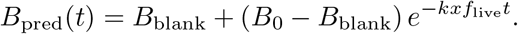

Here *x* is the mean OD used to normalize the corresponding kinetic fit, and *f*_live_ scales the fraction remaining alive. The live–dead mixture measurements were not used to estimate uptake or the release initial condition; however, the fully killed treatment was used to estimate *b*. The remaining live–dead mixture measurements were used to evaluate predictions made using these parameters.

To compare the shapes of the observed and predicted responses across killed-cell fractions, measured B_12_ concentrations were divided by the median measured concentration in the corresponding fully killed treatment. Predicted concentrations were separately divided by the median predicted concentration in the corresponding fully killed treatment. For each biological replicate, the area under the resulting normalized B_12_-versus-killed-fraction curve was calculated by trapezoidal integration across the tested killed-cell fractions. Empirical and predicted area-under-the-curve (AUC) values were then averaged across experiments and biological replicates for each strain. Agreement between empirical and predicted strain-level AUCs was quantified using the mean absolute error (MAE) and Lin’s concordance correlation coefficient.

The statistical significance of the correspondence between empirical and predicted AUCs was assessed using a strain-label permutation test. Predicted AUCs were randomly reassigned among the 23 strains while the empirical AUCs were held fixed, and the MAE was recalculated for each of 1,000 permutations. A one-sided permutation *p*-value was calculated as the proportion of permutations producing an MAE less than or equal to the observed MAE, with one added to both the numerator and denominator.

#### 4.7.2 Supplementary comparison with a quota-limited uptake model

A two-parameter quota model was tested alongside the first-order model in Fig. S12, S13, S14 but not shown in the main figures. Its uptake flux per unit OD was defined as

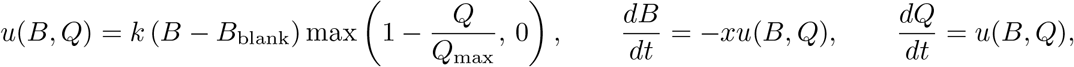

where *Q* is cumulative uptake per unit OD since the start of the assay, *Q*(0) = 0, and *Q*_max_ is the fitted uptake capacity per unit OD 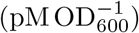. Thus, uptake slows as this capacity is filled; *Q* does not represent a measured initial intracellular B_12_ pool. Uptake time courses used *B*(0) = *B*_blank_ +*B*_0,nominal_. For release time courses, the corresponding uptake parameters were held fixed and the release coefficient was fitted using

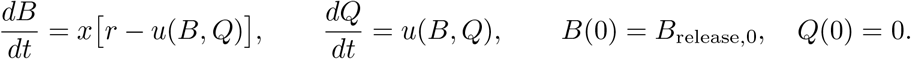

The supplementary first-order comparator reused the fits and predictions from the main analysis, with *u*(*B*) = *k*(*B* − *B*_blank_). Both model structures used the mean initial and final blank-adjusted OD_600_ for the corresponding strain, glucose treatment, experiment and biological replicate, as described above. Only finite, positive OD and B_12_ measurements were used. Parameters minimized the sum of squared residuals between log observed and log predicted concentrations, retaining *B*_blank_ as an additive assay offset. Quota-model parameters were optimized on a log scale. The quota uptake bounds were 10^−8^ ≤ *k* ≤ 10^3^ and max(10^−8^, *σ*_blank,500_*/x*) ≤ *Q*_max_ ≤ 10^6^, in the units defined above, where *σ*_blank,500_ is the standard deviation of B_12_ measurements in blank cultures receiving 500 pM added B_12_. Quota uptake trajectories and live–dead predictions were evaluated using their analytical solution; quota release trajectories were integrated using the adaptive LSODA solver with relative and absolute tolerances of 10^−8^. Numerical accuracy was verified using tolerances of 10^−10^ and checks of nonnegativity, uptake capacity and mass balance.

For the supplementary live–dead comparison (Fig. S13), both model structures predicted uptake following instantaneous release from killed cells. Culture density *x* was the same mean initial and final blank-adjusted OD_600_ used for the corresponding kinetic fit, and *x*_live_ = *xf*_live_. With *b* estimated from the blank-corrected fully killed treatment, predictions used

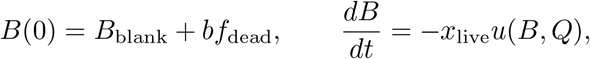

with *Q*(0) = 0 and *dQ/dt* = *u*(*B, Q*) for the quota model, or *u*(*B*) = *k*(*B* − *B*_blank_) for the first-order comparator. These predictions were normalized separately to their corresponding fully killed predictions. In Fig. S12, S13, S14, empirical points show means and standard deviations across experiments and biological replicates; model lines join the means of replicate-specific predictions evaluated at the experimental sampling points. The blank control is displayed but is not fitted in this supplementary comparison.

### 4.8 DNA extraction and genome sequencing

For 91 samples, approximately 5 × 10^9^ cells per sample were preserved in DNA/RNA Shield (Zymo Research) and submitted to Plasmidsaurus (USA) for DNA extraction and standard bacterial whole-genome sequencing between October 2025 and May 2026. For an additional 39 samples, approximately 5 × 10^9^ cells per sample were preserved in DNA/RNA Shield and submitted to Zymo Research for DNA extraction and metagenomic shotgun sequencing in June 2025. For sequencing by Plasmidsaurus, genomic DNA was prepared using an amplification-free library-construction protocol with Oxford Nanopore Technologies (ONT) V14 chemistry and sequenced using ONT R10.4.1 flow cells. Raw signals were basecalled using the Dorado super-accuracy (SUP) model with the default Q10 quality filter. Read quality was further refined by removing the lowest-quality 5% of reads using Filtlong v0.2.1.

### 4.9 Read processing, genome assembly and quality control

Raw Illumina paired-end reads (2 × 151 bp) from 39 samples were adapter-trimmed and quality-filtered using fastp (v1.0.1). Quality reports were generated using FastQC (v0.12.1). Trimmed reads were assembled into contigs using Unicycler (v0.5.1) via the nf-core/bacass pipeline (v2.6.0; Nextflow v26.04.0), which applies an iterative SPAdes-based assembly strategy optimized for short-read bacterial genomes. Assembly quality metrics, including N50, total assembly length, and contig count, were assessed using QUAST (v5.3.0). The resulting assemblies were combined with 223 previously assembled genomes, producing a total of 262 genomes. These assemblies were quality-filtered using a custom Nextflow pipeline (v26.04.0) with CheckM2 (v1.1.0; database: `uniref100.KO.1.dmnd`; completeness ≥ 90%, contamination ≤ 5%) and GUNC (v1.1.0) against the ProGenomes 2.1 database (`pass.GUNC = True`), yielding 231 genomes.

### 4.10 Taxonomic assignment and phylogenomic tree construction

Taxonomic classification was performed using GTDB-Tk (v2.7.2; database release R232) via the `classify wf` workflow. This workflow assigns taxonomy through ANI-based screening against GTDB reference genomes where possible; otherwise, it places genomes into the GTDB reference tree using pplacer (v1.1.alpha19), followed by relative evolutionary divergence (RED)-based assignment. A maximum-likelihood phylogenetic tree was independently inferred using the GTDB-Tk `de novo wf` workflow. The bac120 marker genes were identified in each genome and aligned to the GTDB reference multiple-sequence alignment, after which a tree was inferred using FastTree (v2.2.0). The resulting tree was pruned in R using `ape::keep.tip()` (ape v5.8-1) to retain selected genomes with measured phenotypes.

### 4.11 Genome annotation and KEGG ortholog assignment

Quality-filtered genomes were annotated using Bakta (v1.11.4; database v6.0), which predicts protein-coding sequences (CDS), rRNA, tRNA, and other non-coding features. Predicted protein sequences were assigned KEGG Orthology (KO) terms using KofamScan (v1.3.0) against KEGG HMM profiles (release 2026-04-28) and KO list (release 2026-04-27); per-genome KO assignments were aggregated into a genome × KO count matrix and binary matrix for downstream functional analysis.

### 4.12 Random forest prediction of B_12_ traits

Binary random forest classifiers were trained using genome-wide KEGG Orthology (KO) presence–absence profiles. Four classification tasks were considered. Across all strains, models predicted B_12_ synthesis (total B_12_ ≥ 50) and B_12_ uptake (≥ 250 pM). After restricting the dataset to B_12_ synthesizers (total B_12_ *>* 50), separate models predicted B_12_ uptake (≥ 250 pM) and extracellular B_12_ sharing (EC-B12 fraction ≥ 0.05). Feature filtering was performed independently within each training set to prevent information leakage. KOs present in fewer than 2% or more than 98% of training genomes, or with no variation, were excluded. A single random forest was then fitted using all remaining KOs, without subsequent feature selection or model refitting. Models were fitted using the `ranger` R package with 1,000 trees, 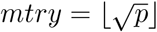, a minimum terminal-node size of one, and a 63.2% sample fraction without replacement. Other parameters were left at their default values.

Predictive performance was evaluated using two test-set designs. Stratified random validation used seven non-overlapping folds for all strains and six for synthesizers, matching the number of order blocks. For phylogenetically structured splits, GTDB orders representing at least 5% of the relevant dataset were held out separately, while smaller orders were pooled to produce a test set meeting the same minimum size. Each strain therefore occurred in only one order-block test set.

For each split, null performance was estimated from 1,000 sets of Bernoulli predictions sampled according to the class prevalence in the corresponding training set. Classifier accuracy was compared with the mean null accuracy for each matched split. Evidence that classifier performance exceeded null performance was assessed using a one-sided paired *t*-test across splits.

### 4.13 Genome-wide association (GWA) analysis with pyseer

To conduct the GWA analysis, we used the pyseer software developed by Lees et al. (2018). Pyseer requires a binary input matrix, so we first binarized the KO count values to be 1 if the count exceeded 0, and 0 otherwise. Next, groups of KO count columns that were identical (perfectly correlated) were collapsed into single columns. When presenting the results, all KOs in the same group were assigned the same corresponding output. Binarized KO counts with a minor allele frequency below 0.01 were filtered out, and the remaining counts were standardized by subtracting the KO-specific mean and dividing by the KO-specific standard deviation. We use *X* to denote the resulting matrix of transformed KO counts.

For the phenotypes log 10 total B_12_ gm and B_12_ mean uptake, there were 203 and 201 samples with matching genomes, respectively. For both phenotypes, we fitted pyseer linear mixed models (FastLMM) using two choices of kinship matrix: the *XX*^*T*^ matrix and a kinship matrix derived from the estimated and subsequently pruned phylogenetic tree. In R (v4.5.2), tree pruning was performed using the `keep.tip` function, and the phylogenetic kinship matrix was derived using the `vcv.phylo` function. Both functions are implemented in the `ape` package (v5.8-1). Pyseer ran without triggering any quality control flags and produced likelihood-ratio-test *p*-values, which were then corrected for multiple comparisons using the Benjamini–Hochberg and Bonferroni procedures.

### 4.14 Statistical analysis and data visualization

Statistical tests, false discovery rate control, and definitions of error bars are provided in the main text and/or figure legends. No data were excluded from analysis unless they did not meet predefined quality criteria. Statistical analyses and data visualization were performed in R version 4.5.1.

### 4.15 Data and code availability

Data supporting this study are provided in the article and its Supplementary Information. Previously published genome sequences are available under NCBI BioProjects PRJNA660495, PRJNA940744, PRJNA414740, and PRJNA478695. Genome sequences generated in this study have been submitted to NCBI under BioProject PRJNA1529823. Code used for data processing, analysis, and figure generation is available at https://github.com/freddybunbury1/Bacterial_vitamin_sharing.

## 5 Supplementary figures

**Fig. S1.**
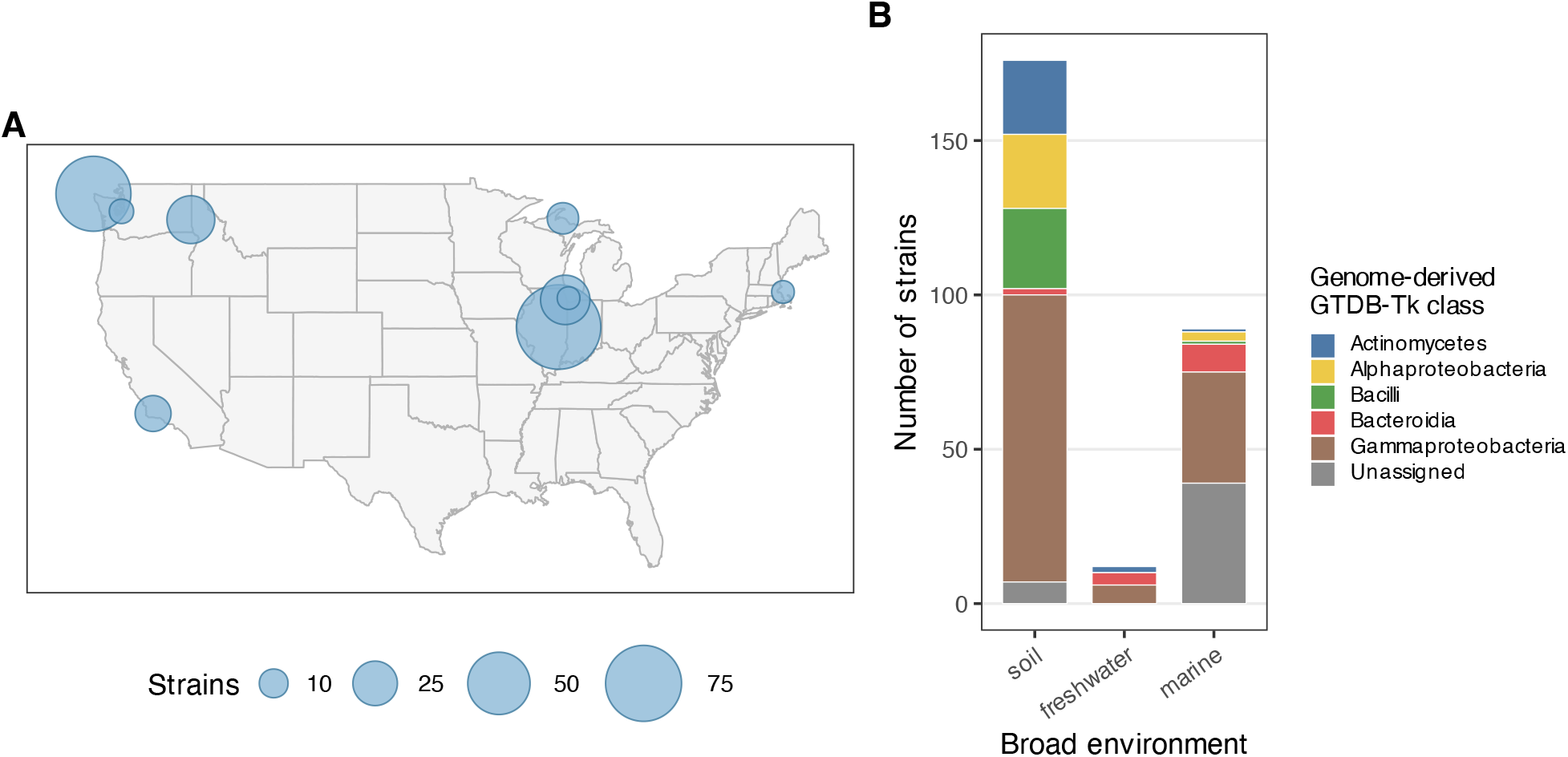
Geographic, environmental, and taxonomic distribution of the bacterial strain collection. (A) Sampling-location cluster centroids across the continental United States. Sampling coordinates were grouped into ten reproducible clusters; bubble area is proportional to the number of strains assigned to each cluster. (B) Numbers of strains isolated from soil, freshwater and marine environments, stacked by genome-derived GTDB-Tk class. Grey segments denote strains without an assigned GTDB-Tk class.

**Fig. S2.**
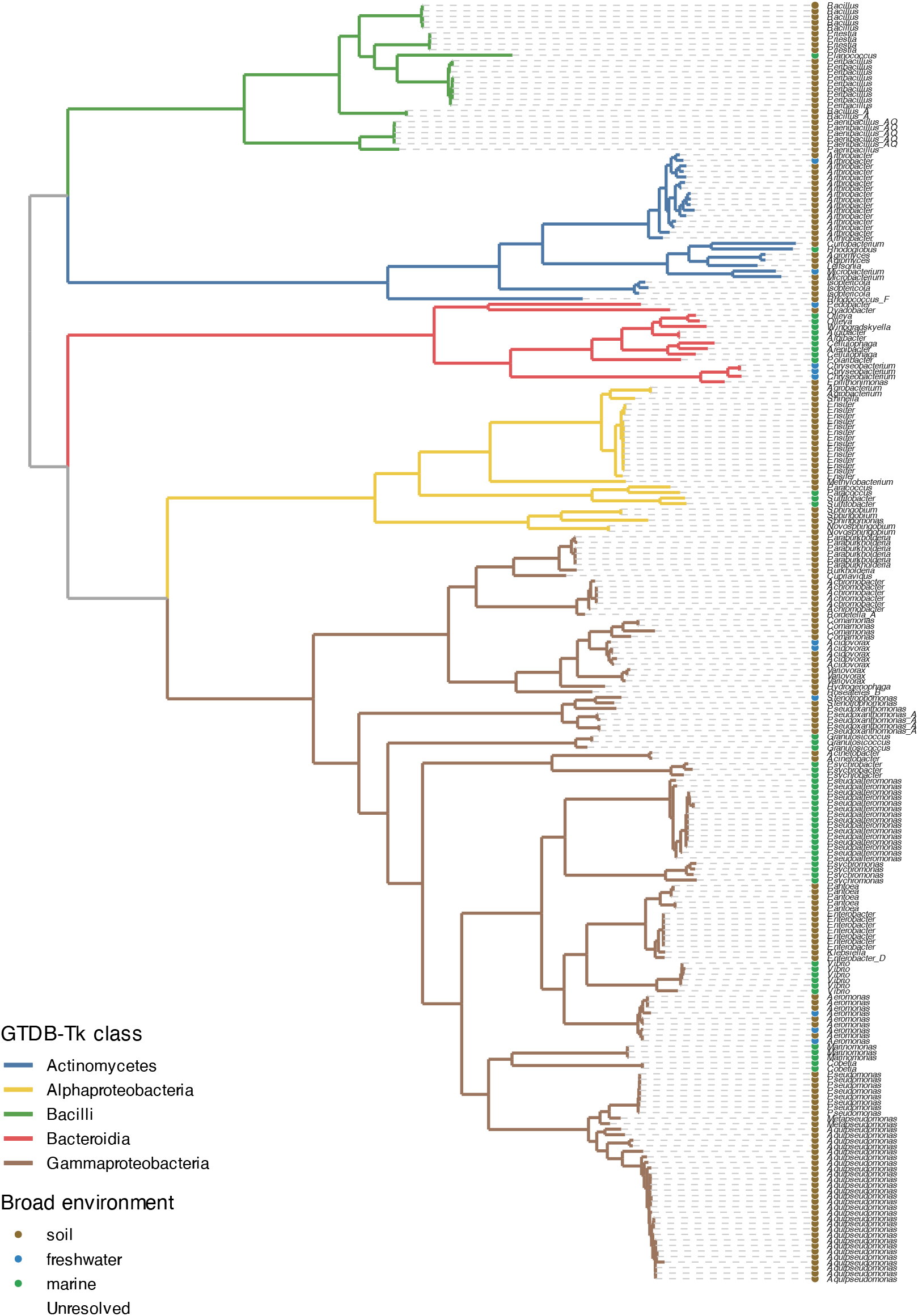
Phylogenetic relationships among the 231 bacterial genomes retained for phylogenomic analysis. The tree was inferred from aligned bac120 marker genes using FastTree within the GTDB-Tk de novo wf workflow and rooted at the best-supported split between Bacillati and Pseudomonadati. Tip labels indicate GTDB-Tk genus assignments, tip-point colours indicate the broad environments from which the isolates were collected, and branch colours indicate GTDB-Tk class.

**Fig. S3.**
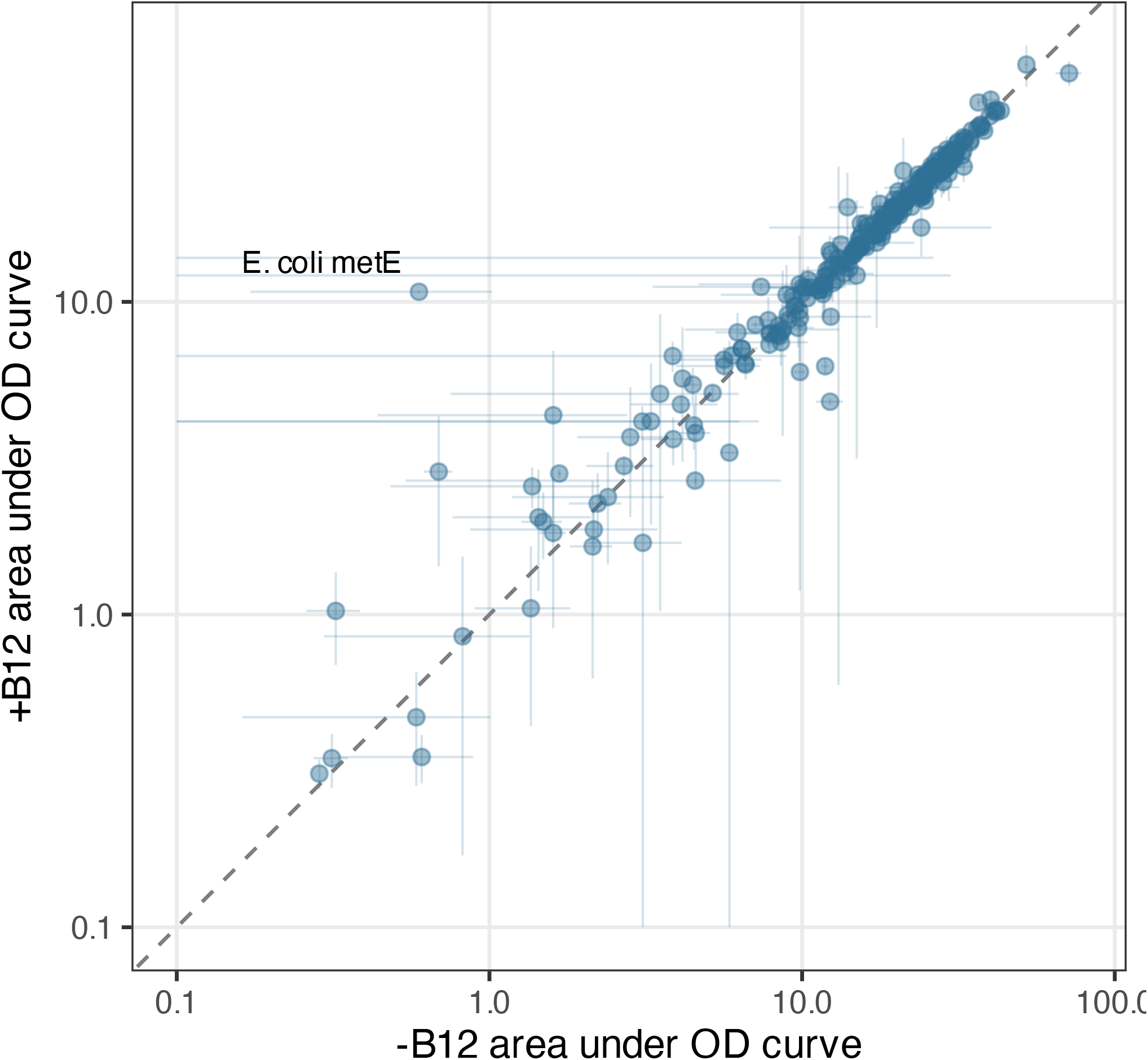
Growth of 274 bacterial collection isolates and controls in Tm-mod medium with (y-axis) or without (x-axis) 10nM added B_12_. Growth was assessed as the area under the optical density at 600nm curve over 57 hours with measurements every 93 minutes. The B_12_- or methionine-dependent *E. coli metE* reporter strain was the only strain that showed substantial deviation from equal growth with and without B_12_ (1:1 diagonal dashed line).

**Fig. S4.**
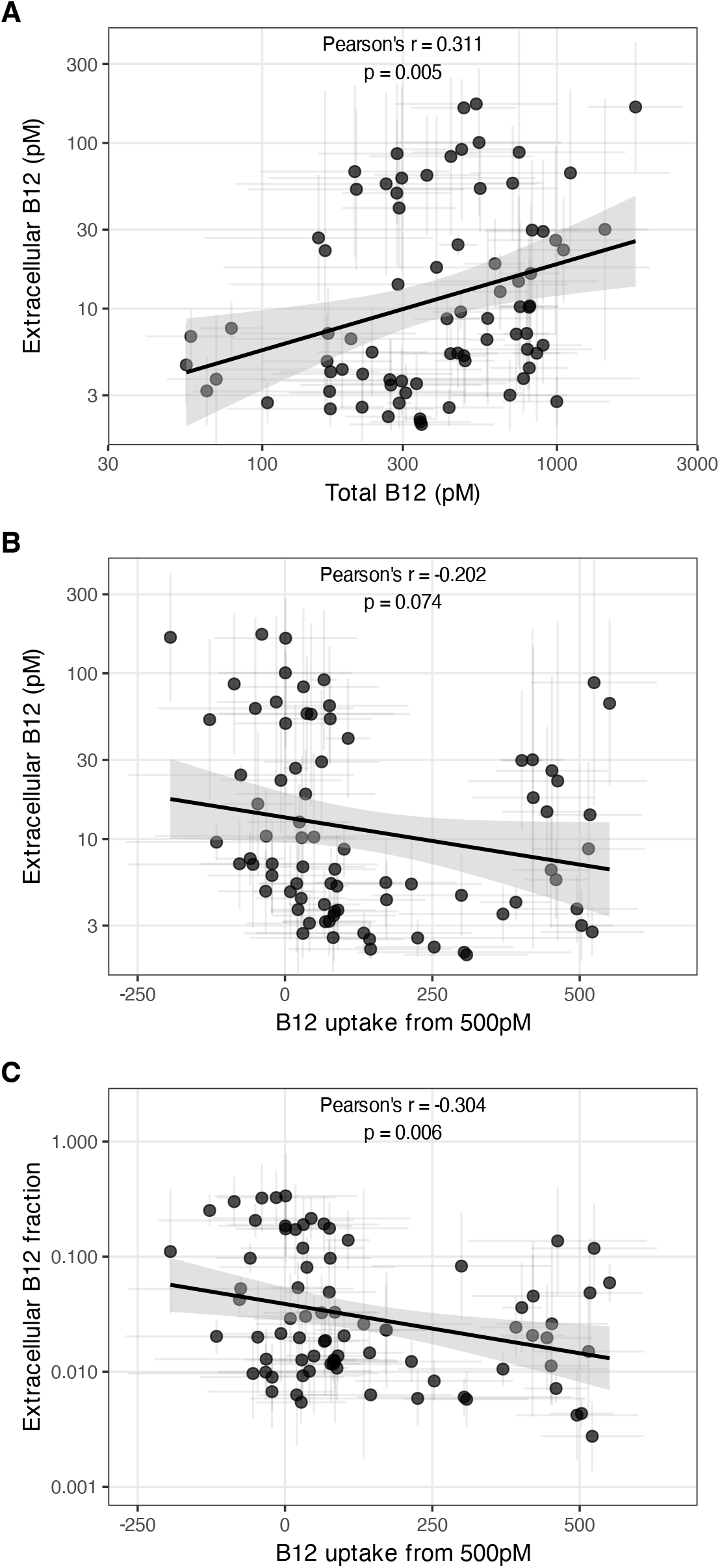
Relationships among total B_12_, extracellular B_12_, B_12_ uptake and the extracellular B_12_ fraction among B_12_ synthesizers (*>* 50 pM total B_12_). (A) Extracellular versus total B_12_. (B) Extracellular B_12_ versus uptake from an initial concentration of 500 pM. (C) Extracellular B_12_ fraction versus uptake. Points represent individual strains. Error bars for total B_12_, extracellular B_12_ and extracellular B_12_ fraction show geometric standard deviations, whereas uptake error bars show arithmetic standard deviations, calculated from up to four measurements comprising two biological replicates at two time points. Lines show linear-regression fits on the plotted scales, and shaded regions show 95% confidence intervals. Pearson’s *r* and the *P* -value for the null hypothesis *r* = 0 are shown at the top of each panel.

**Fig. S5.**
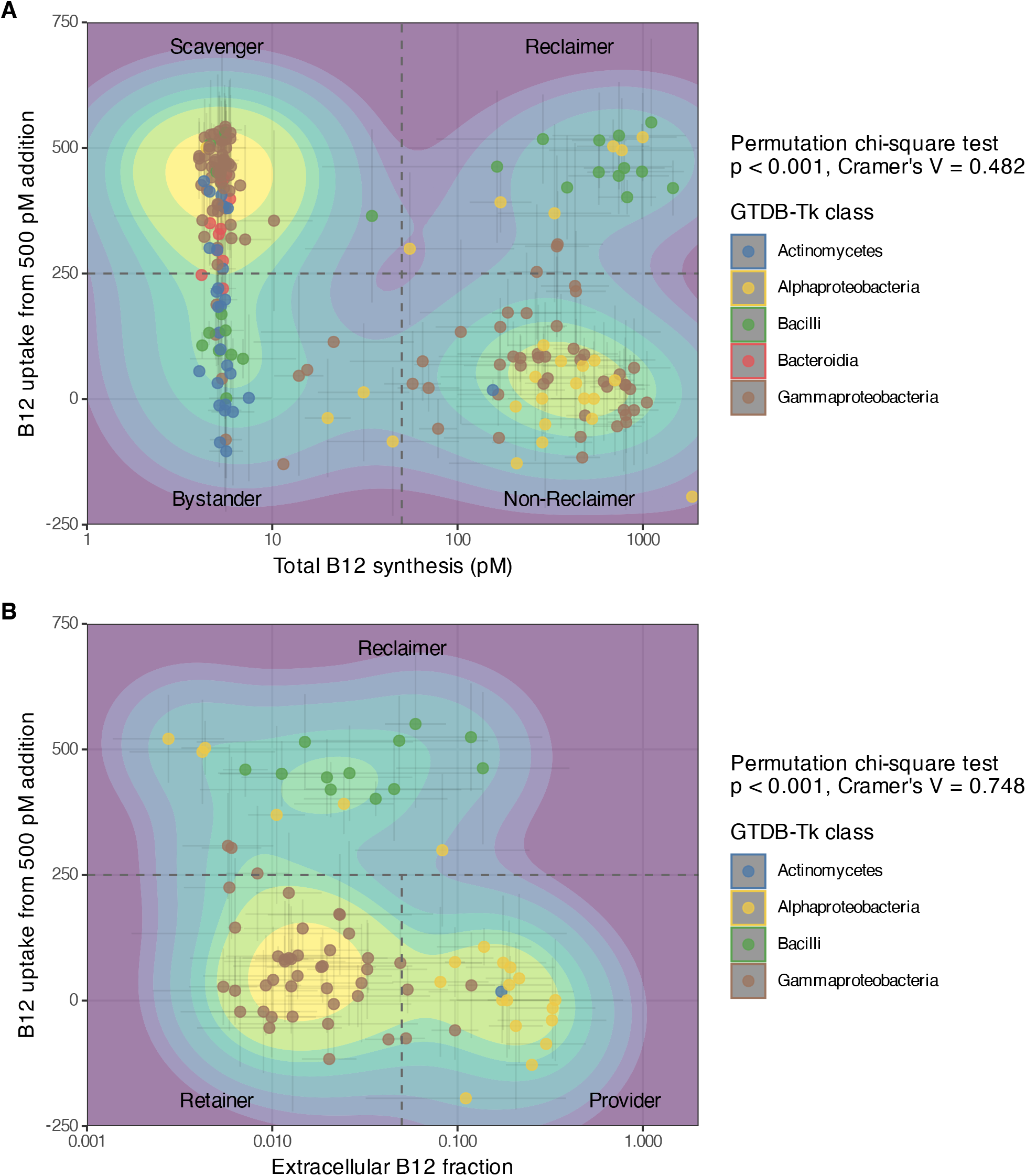
B_12_ trait space across bacterial taxonomic classes. (A) Total B_12_ versus uptake from an initial concentration of 500 pM for all strains with sufficient measurements. Dashed lines indicate the thresholds used to distinguish bystanders, scavengers, reclaimers, and non-reclaimers: 50 pM total B_12_ and 250 pM uptake. Trait-group membership was associated with GTDB-Tk class (permutation chi-square test with 9,999 permutations: *p <* 0.001, Cramér’s *V* = 0.482). (B) Extra-cellular B_12_ fraction versus uptake among B_12_ synthesizers (*>* 50 pM total B_12_). Dashed lines indicate the thresholds used to distinguish retainers, providers, and reclaimers: an extracellular B_12_ fraction of 0.05 and uptake of 250 pM. Trait-group membership was associated with GTDB-Tk class (permutation chi-square test with 9,999 permutations: *p <* 0.001, Cramér’s *V* = 0.748). Points represent individual strains and are coloured by GTDB-Tk class; background shading represents normalized two-dimensional point density. Strains without an assigned GTDB-Tk class are shown but were excluded from the contingency-table analyses. Error bars for total B_12_ and extracellular B_12_ fraction show geometric standard deviations, whereas uptake error bars show arithmetic standard deviations, calculated from up to four measurements comprising two biological replicates at two time points.

**Fig. S6.**
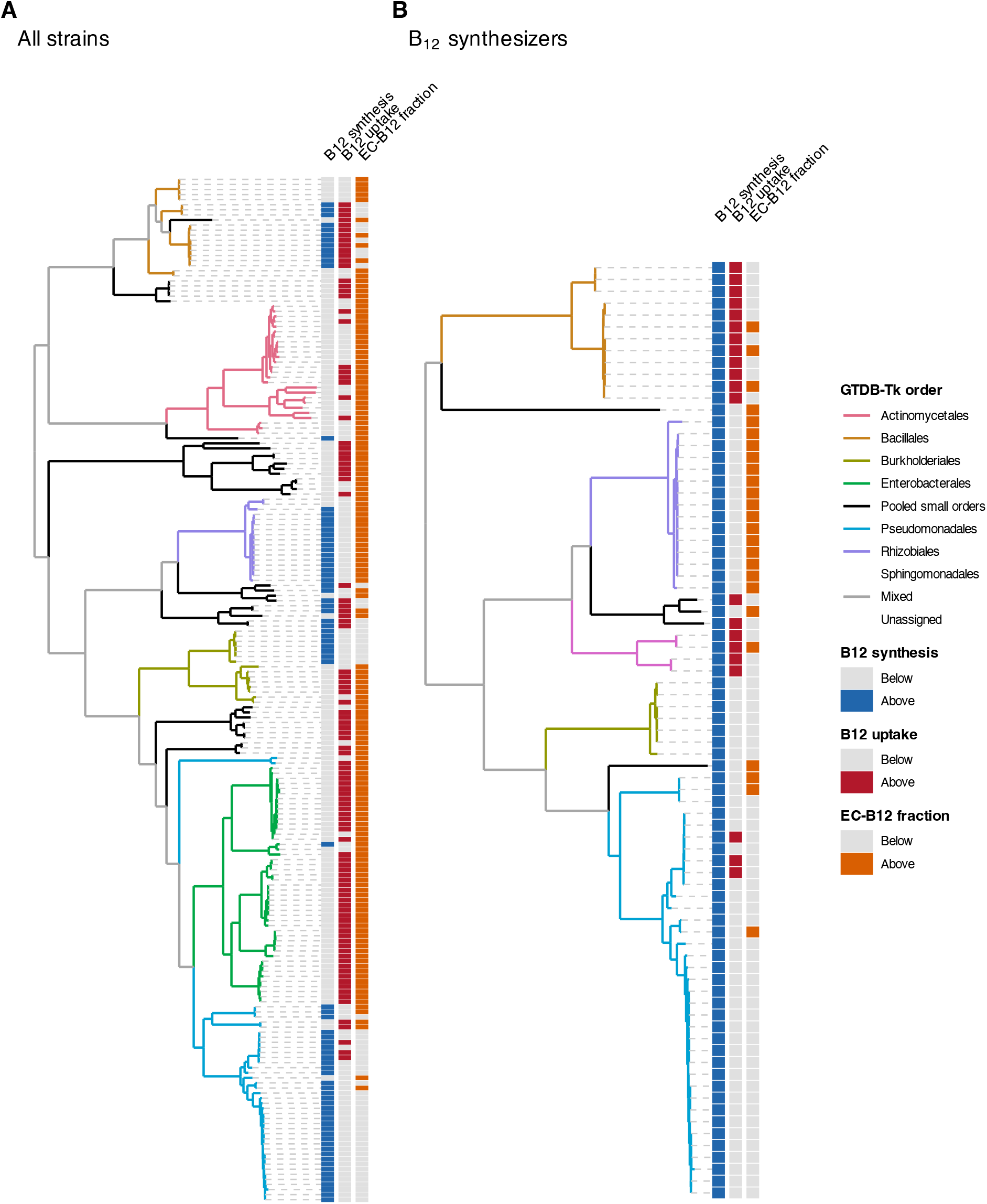
Phylogenetic distribution of binary B_12_ phenotypes. (A) Binary traits across strains with available phenotype data. (B) The corresponding tree restricted to B_12_-synthesizing strains with total *B*_12_ *>* 50 pM and complete uptake and extracellular-fraction measurements. Traits were classified as above the indicated threshold when total *B*_12_ ≥ 50 pM, mean B_12_ uptake ≥ 250 pM, or the extracellular B_12_ fraction ≥ 0.05; all lower values were classified as below the threshold. Both trees were generated by pruning the GTDB-Tk phylogeny shown in Figure S2 to retain the relevant strains. Branch colours denote GTDB order. Orders containing at least 5% of strains in the respective analysis set were represented separately, whereas smaller orders were combined into the pooled smaller-orders group.

**Fig. S7.**
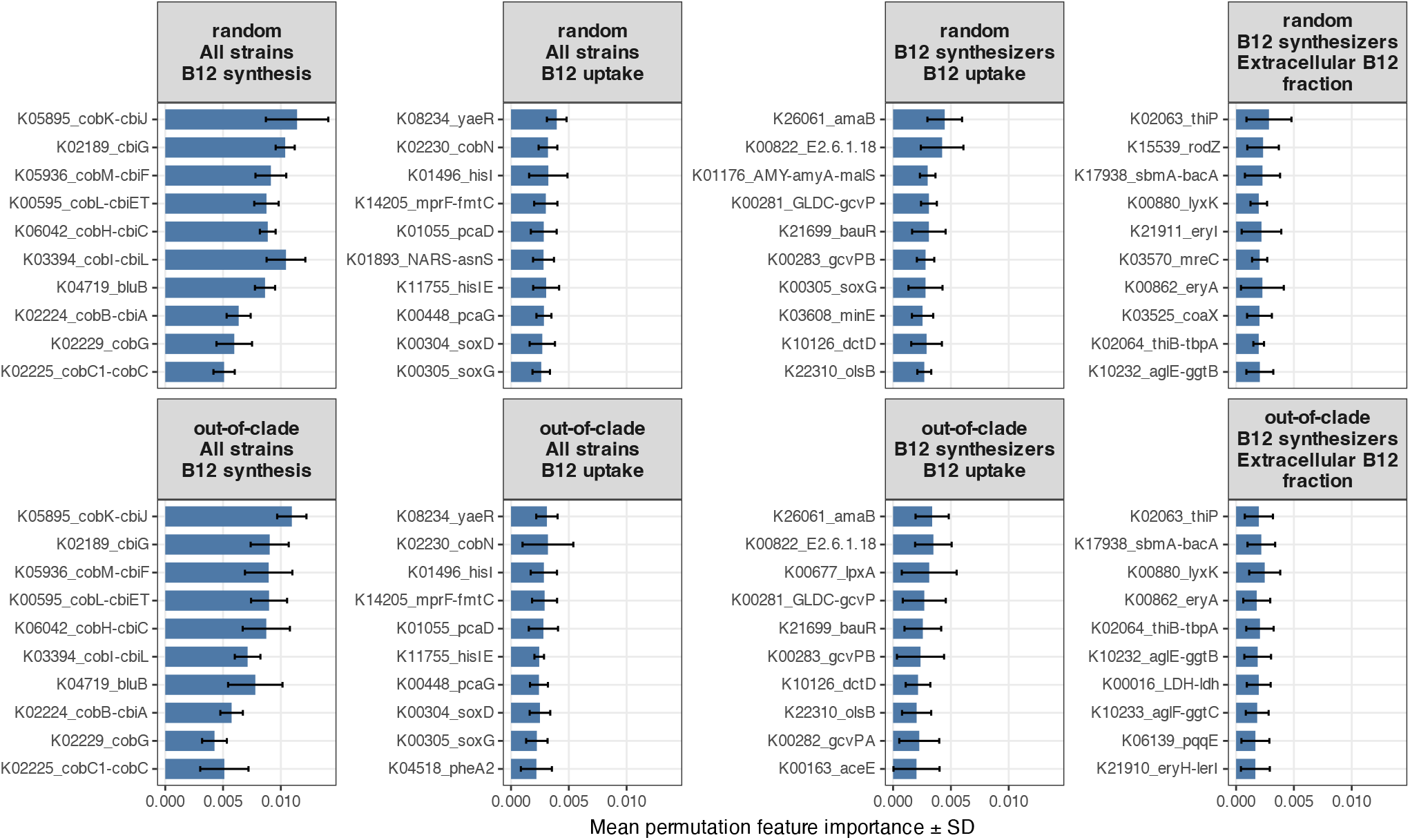
Most informative genomic features for random-forest classification of B_12_ phenotypes. Mean permutation feature importance is shown for the ten most informative features under random stratified and out-of-clade train–test splits. Models were fitted either to all strains or to B_12_ synthesizers with total *B*_12_ *>* 50 pM, as indicated. Binary phenotypes were classified as above the threshold when total *B*_12_ ≥ 50 pM, mean B_12_ uptake ≥ 250 pM, or the extracellular B_12_ fraction ≥ 0.05; lower values were classified as below the threshold. Error bars represent the standard deviation in permutation importance among model fits.

**Fig. S8.**
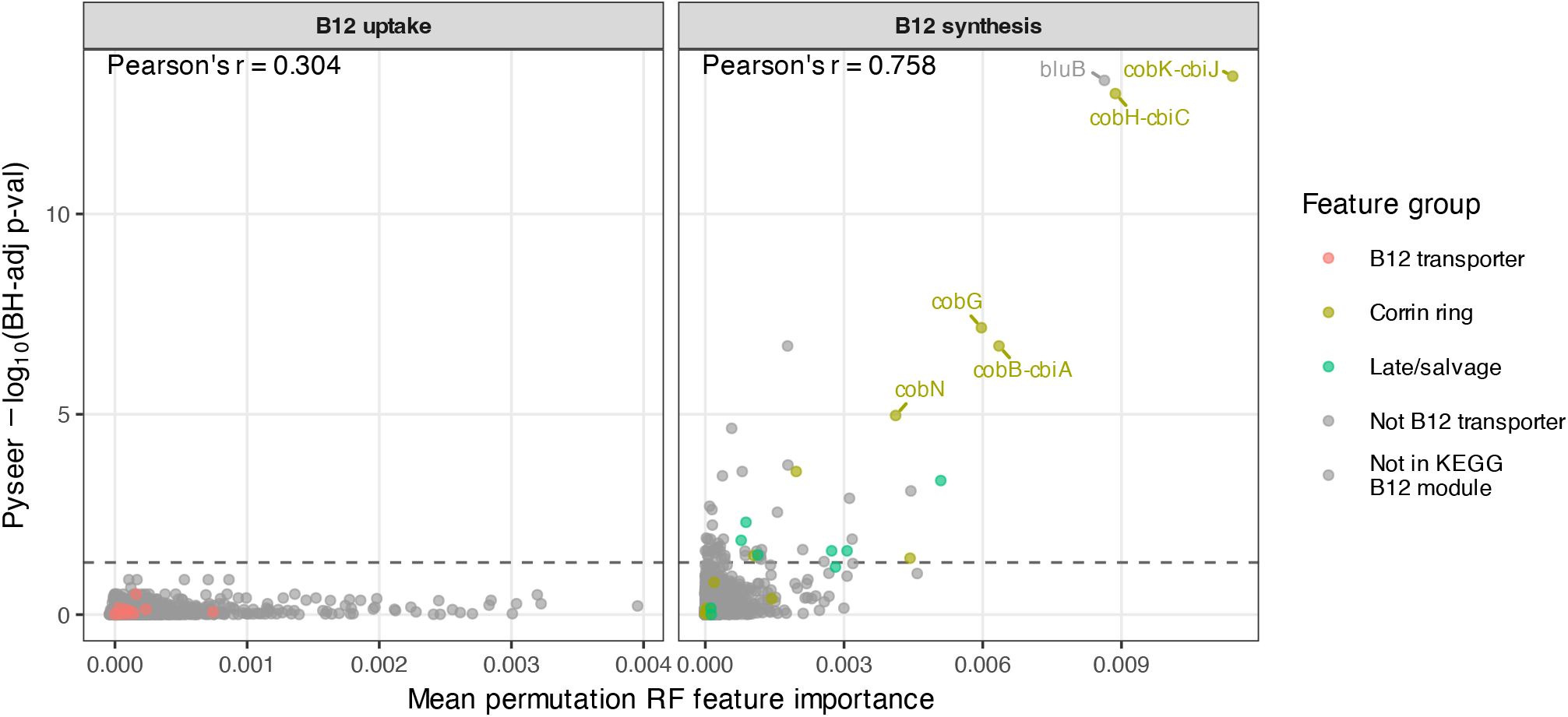
Comparison of random-forest permutation feature importance with population-structure-adjusted Pyseer association evidence for total B_12_ and uptake phenotypes. Pyseer evidence is shown as − log_10_(*P*_BH_), where the likelihood-ratio-test *P* -values were adjusted across all tested KOs for each phenotype using the Benjamini–Hochberg procedure. The 11 candidate B_12_-transporter KOs were K16092 (*btuB*), K06858 (*btuF*), K25034 (*btuF*), K06073 (*btuC*), K25027 (*btuC*), K06074 (*btuD*), K25028 (*btuD*), K02471 (*bacA/bclA*), K16927 (*cbrT*), K01552 (*ecfA*) and K16785 (*ecfT*). Genes assigned to the corrin-ring category were members of the KEGG anaerobic (M00924) or aerobic (M00925) cobalamin-biosynthesis modules; genes in the late/salvage category were members of the KEGG late cobalamin-completion module (M00122). Although *bluB* (K04719) is not annotated as a member of these KEGG B_12_-biosynthesis modules, it synthesizes the lower ligand 5,6-dimethylbenzimidazole. Labelled points identify selected annotated genes; grey points denote KOs not assigned to the candidate B_12_-transporter set or the specified KEGG B_12_ modules. Pearson’s *r* reports the correlation between permutation importance and Pyseer association evidence within each phenotype.

**Fig. S9.**
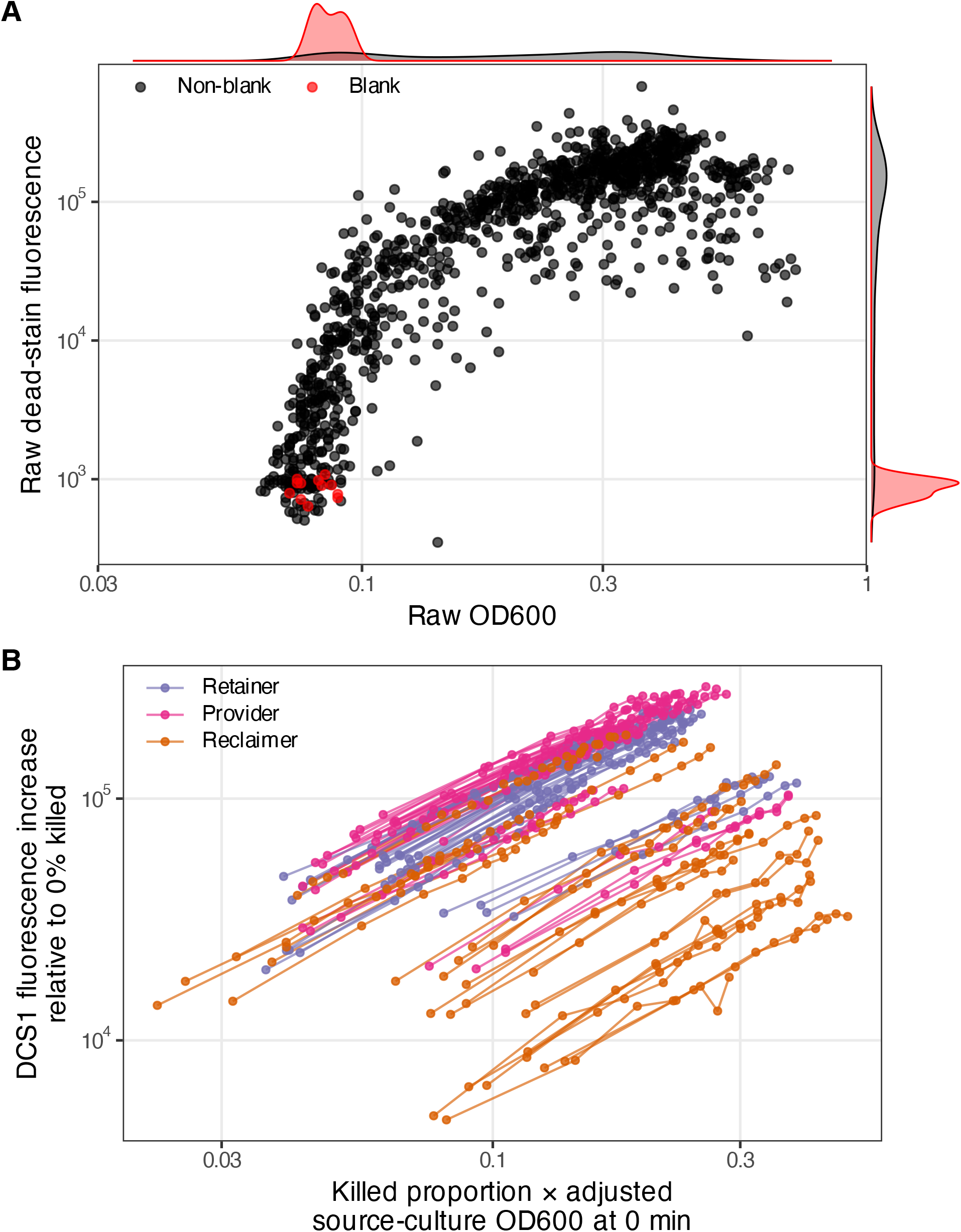
Validation of dead-cell fluorescence against OD_600_ and controlled live–dead mixtures. Fluorescence was measured using the dead-cell-specific dye Nuclear Green DCS1 (AAT Bioquest; catalogue no. 50-224-9741). (A) Raw OD_600_ before killing versus raw fluorescence after heat killing (95 °C for 30 min) for samples from the bacterial B_12_ trait survey in Fig. 1. Points represent individual culture samples; non-blank samples are black and blanks are red. Marginal densities show the log-transformed OD_600_ and fluorescence distributions. (B) Cultures of 23 selected B_12_-producing strains were divided into untreated live and heat-killed fractions (95 °C for 30 min), then mixed to produce cultures containing 0, 28.5, 50.5, 66.5, 79, 88, 95 or 100% killed cells. The x-axis shows the experimentally imposed killed-cell fraction multiplied by the blank-adjusted OD of the corresponding source culture at time zero. The y-axis shows the increase in DCS1 fluorescence relative to the matched 0%-killed sample from the same strain and replicate. Lines connect measurements within these matched groups; colours indicate retainer, provider and reclaimer trait groups.

**Fig. S10.**
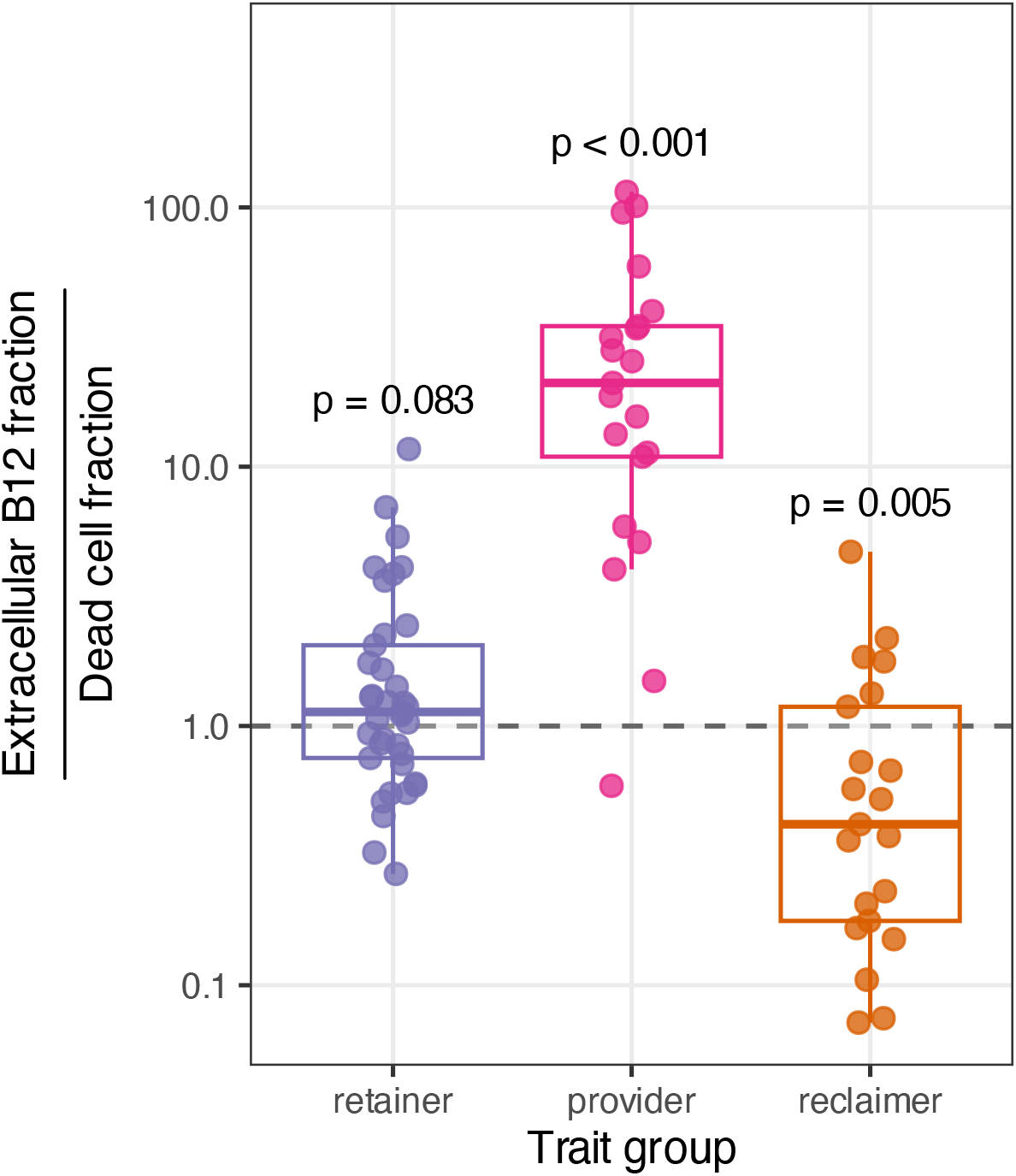
Extracellular B_12_ fraction normalized by dead-cell fraction across B_12_ trait groups. Points represent strain-level observations, and boxplots summarize the corresponding trait-group distributions. The dashed horizontal line denotes a ratio of one. Within each trait group, the annotated *P* -value was calculated using a two-sided one-sample Student’s *t*-test of the log_10_-transformed ratios against zero, corresponding to testing whether the geometric mean untransformed ratio differs from one.

**Fig. S11.**
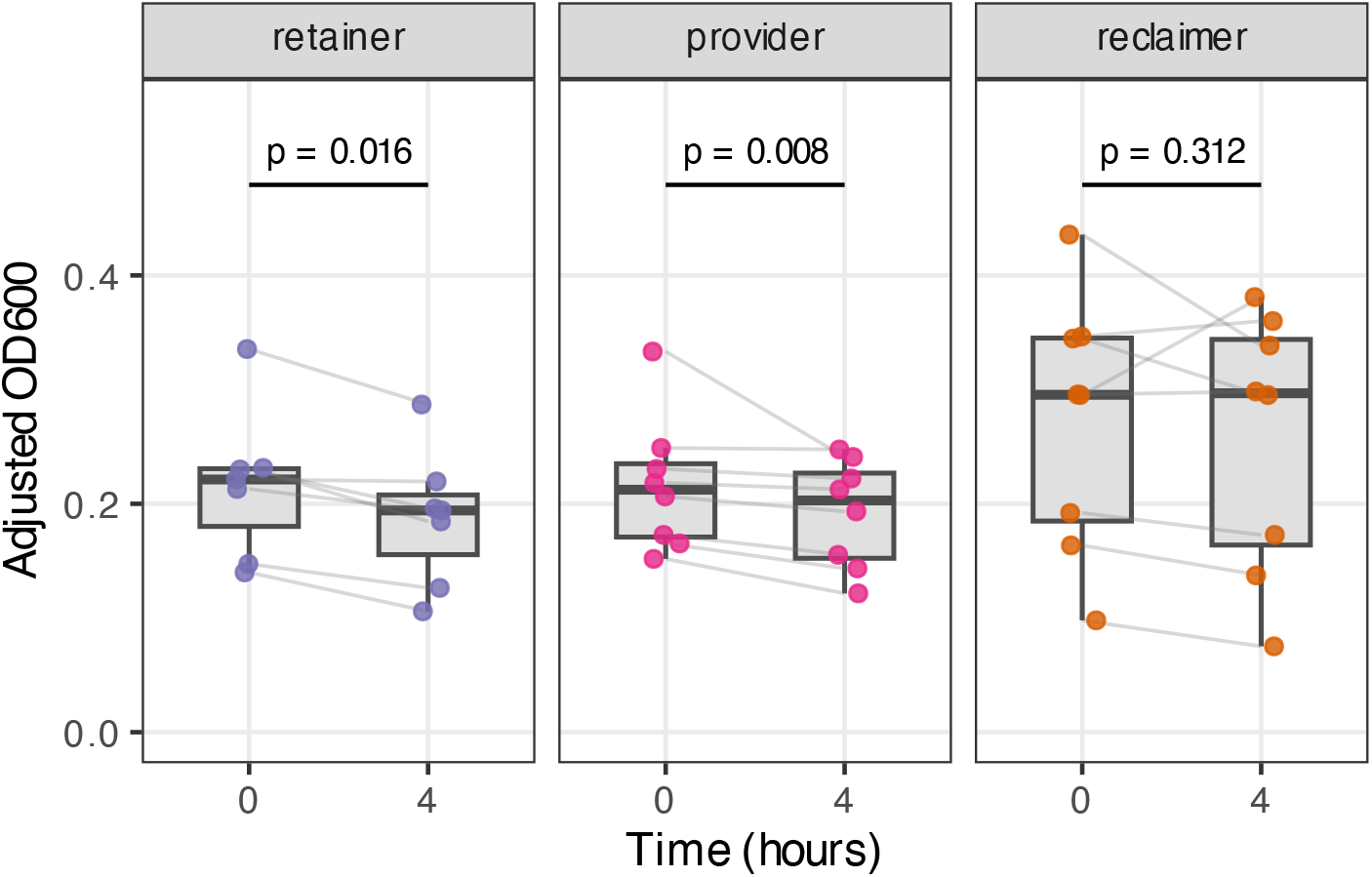
Initial and final culture density during the B_12_ uptake assay without added glucose. OD_600_ was measured in the source cultures before B_12_ addition and after four hours in the corresponding wells supplied with 500 pM B_12_. Measurements were blank-subtracted. Points represent strain-level means across four matched cultures, lines connect the same strain, and boxplots summarize the corresponding trait-group distributions. Within each trait group, the annotated *P* -value was calculated using a two-sided paired Wilcoxon signed-rank test comparing initial and final strain-level OD_600_.

**Fig. S12.**
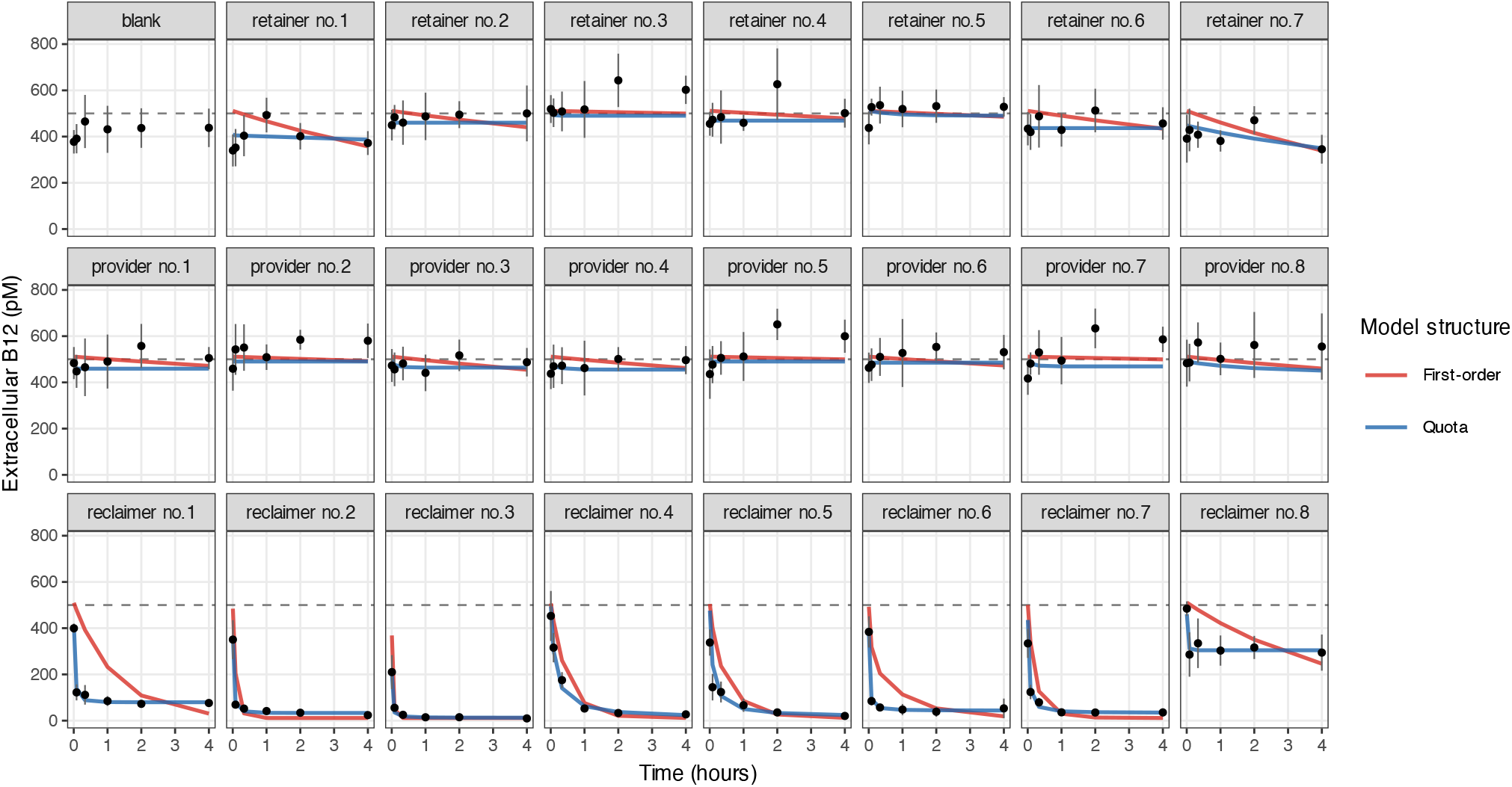
Empirical extracellular B_12_ measurements and fitted uptake-model predictions in the absence of added glucose. First-order and quota-limited uptake models were fitted to log-transformed extracellular B_12_ concentrations. The concentration at *t* = 0, *B*_0_, was fixed as the sum of the experimentally added B_12_ concentration and the common blank offset, *B*_blank_. The OD-specific uptake coefficient *k* was estimated from the subsequent time course; the quota model additionally fitted an uptake capacity *Q*_max_, with uptake slowing as this capacity was filled. Points show empirical measurements; red and blue lines show first-order and quota-model predictions, respectively. Facets show the blank and individual strains ordered within retainer, provider and reclaimer trait groups.

**Fig. S13.**
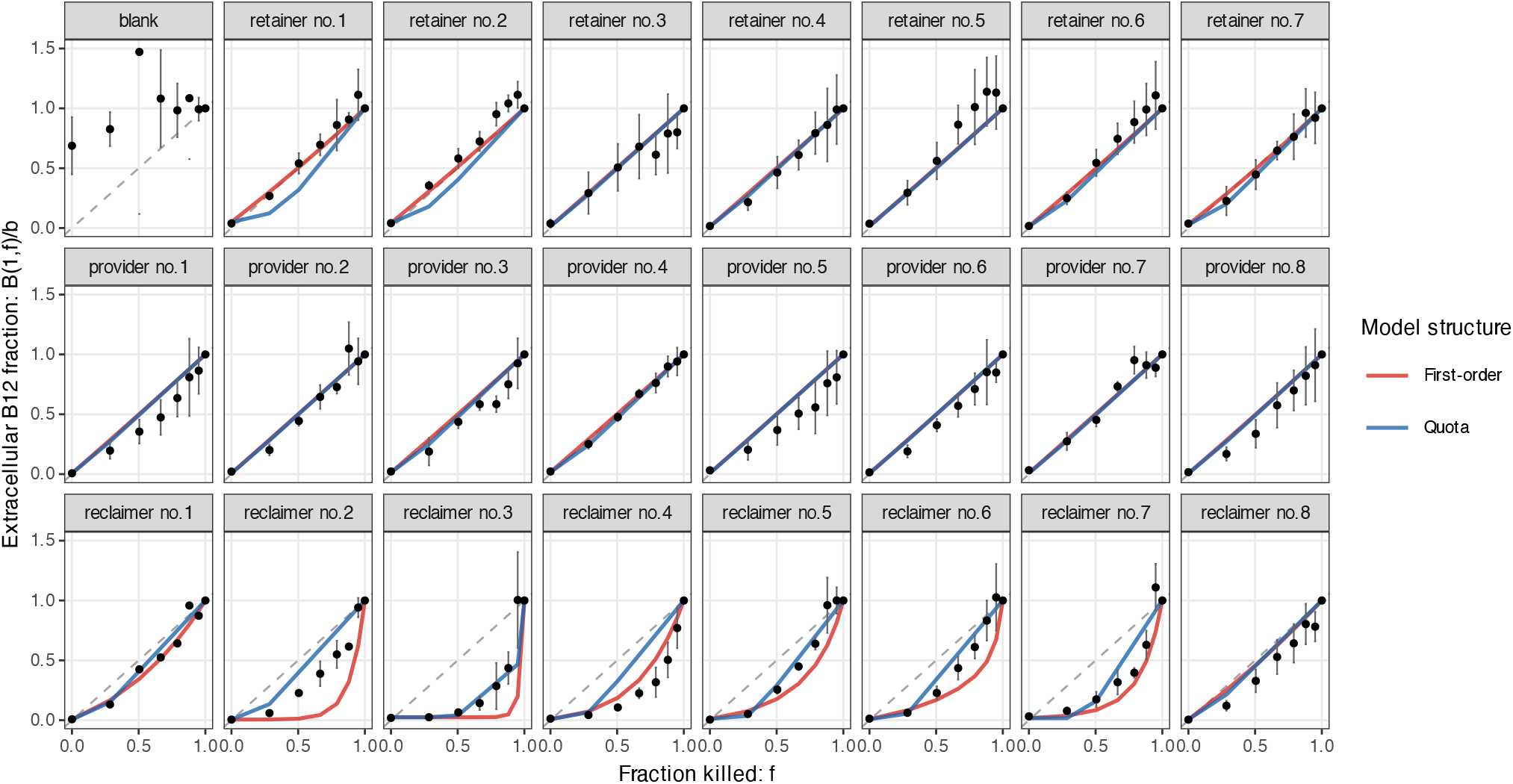
Prediction of extracellular B_12_ availability in controlled live–dead cell mixtures using independently estimated uptake parameters. For each strain, the first-order and quota-limited models assume that killed cells instantaneously release intracellular B_12_ in proportion to the dead-cell fraction *f*, while the surviving fraction 1 − *f* recaptures extracellular B_12_ according to the uptake parameters estimated from the corresponding uptake experiment (Fig. S12). Empirical and predicted extracellular B_12_ concentrations after 1 h were normalized to their corresponding empirical and predicted concentrations in the fully heat-killed culture (*f* = 1). Points show normalized empirical measurements; red and blue lines show first-order and quota-model predictions, respectively. The dashed diagonal indicates equality between the extracellular B_12_ fraction and the fraction killed. Facets show the blank and individual strains ordered within retainer, provider and reclaimer trait groups.

**Fig. S14.**
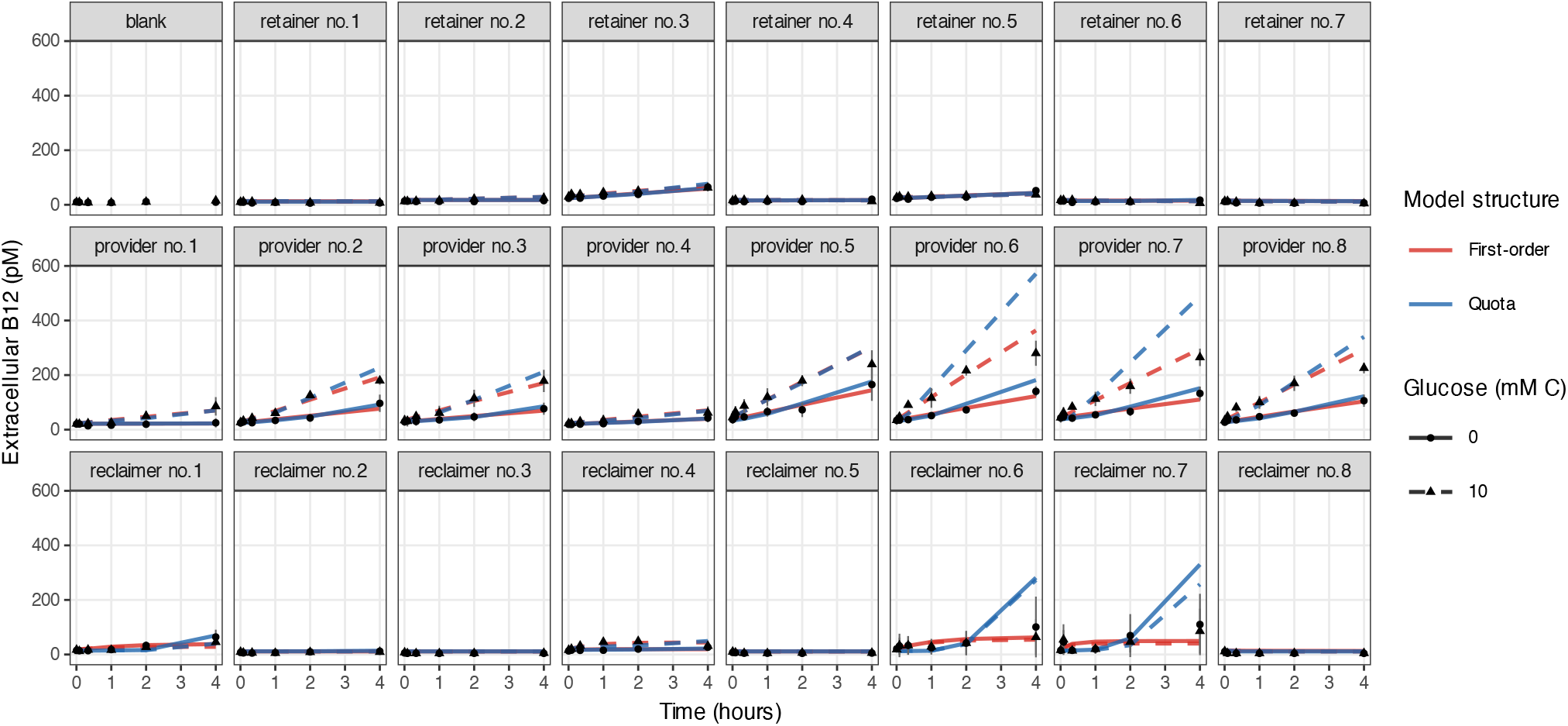
Empirical extracellular B_12_ measurements and release-model predictions with or without added glucose. Glucose was supplied at 10 mM carbon atoms. Predictions were generated using the first-order and quota-limited models described in Methods. For each model, the corresponding uptake parameters were estimated independently from the uptake experiment and held fixed, while the OD-specific release-rate coefficient *r* was fitted. The quota model additionally limits cumulative uptake through the fitted capacity *Q*_max_. Both models account for culture density using the mean initial and final blank-adjusted OD_600_ and retain the common blank offset. Points show empirical measurements; red and blue lines show first-order and quota-model predictions, respectively. Dashed lines and triangles denote cultures receiving glucose; solid lines and circles denote cultures without added glucose. Facets show the blank and individual strains ordered within retainer, provider and reclaimer trait groups.

**Fig. S15.**
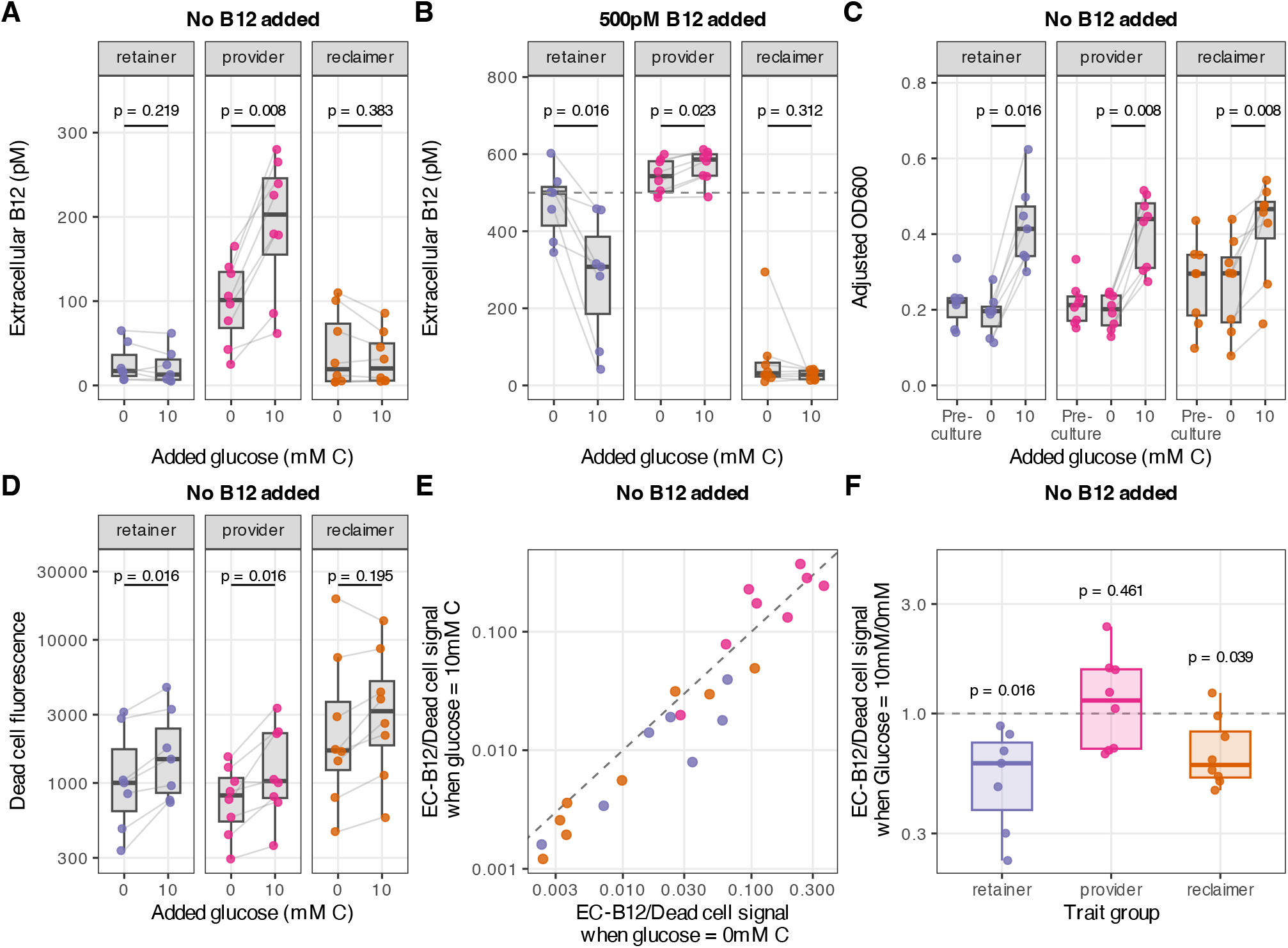
Effects of added glucose on extracellular B_12_, culture density and dead-cell signal after 4 h of culturing. Glucose treatments were 0 or 10 mM carbon atoms (1.67 mM glucose). All panels except panel B show cultures without added B_12_. (A) Extracellular B_12_ without added B_12_. (B) Extracellular B_12_ following the addition of 500 pM B_12_; the dashed horizontal line indicates the added concentration. (C) Blank-adjusted OD_600_ before experimental setup (Pre-culture) and after 4 h without added B_12_, with or without glucose. (D) Dead-cell fluorescence without added B_12_, shown on a logarithmic scale. In panels A–D, points represent strain-level means, grey lines connect the same strain between glucose treatments and boxplots summarize the distributions. Facets separate retainers, providers and reclaimers. Annotated *P* -values are from two-sided exact paired Wilcoxon signed-rank tests comparing the two glucose treatments within each trait group. (E) Strain-level extracellular B_12_ (from panel A) normalized by dead-cell signal (from panel D) under 10 versus 0 mM glucose carbon, with both axes shown on logarithmic scales. The dashed diagonal denotes equal normalized extracellular B_12_ under the two treatments. (F) The corresponding within-strain ratio between the 10 and 0 mM glucose-carbon treatments, summarized by trait group; the dashed horizontal line at one denotes no glucose-associated change.

**Fig. S16.**
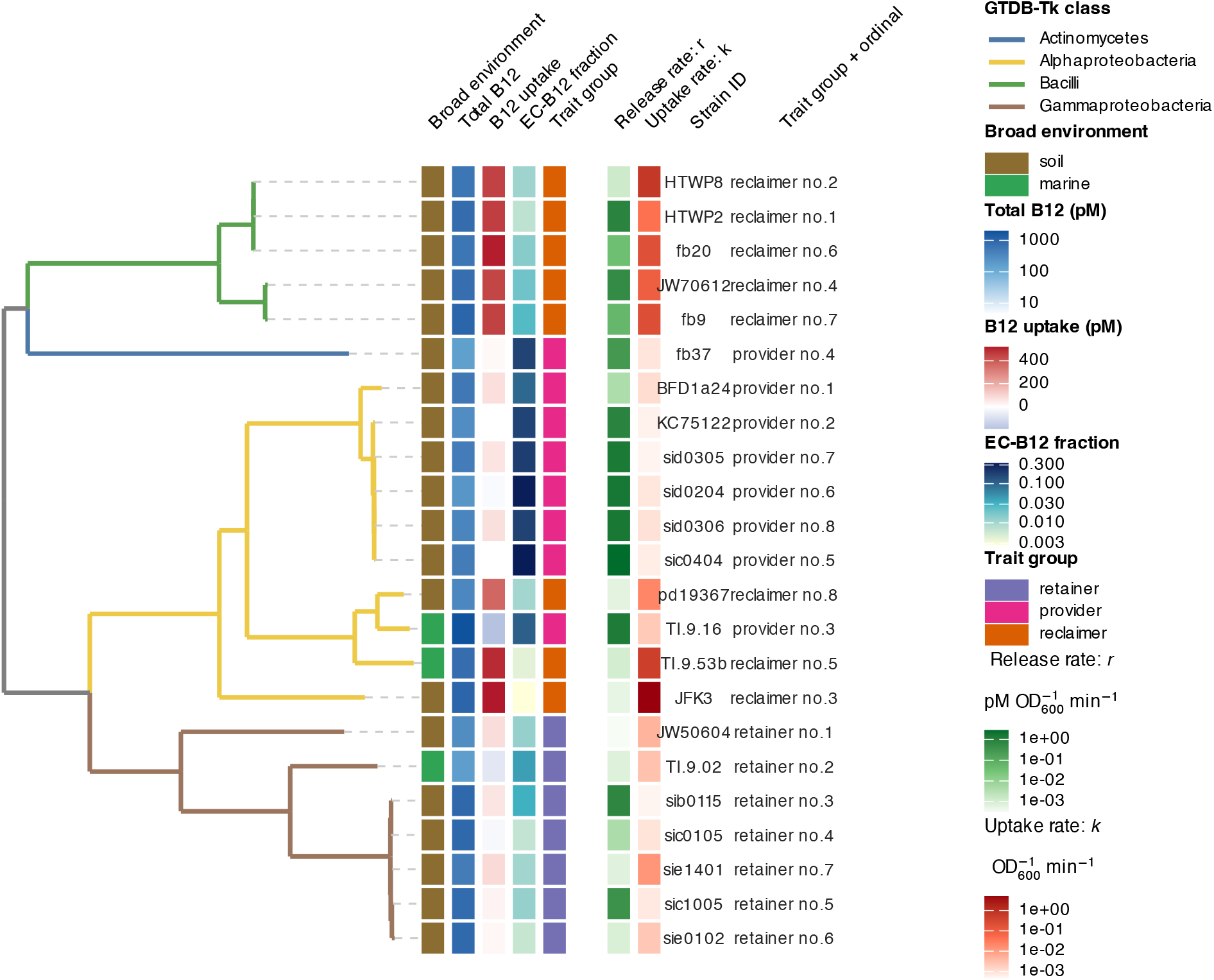
Phylogenetic distribution of B_12_-related traits and first-order release and uptake rates among the 23 strains selected for dynamic measurements. The pruned GTDB-Tk phylogeny retains branch lengths, with branch colours indicating taxonomic class and dashed grey lines linking terminal branches to the corresponding data rows. Columns show, from left to right, broad environmental origin; total B_12_; B_12_ uptake; the extracellular B_12_ fraction (EC-B_12_ fraction); trait group; the OD-specific release-rate coefficient *r*; the uptake-rate coefficient *k*; strain identity; and the strain’s trait-group ordinal. Total B_12_, EC-B_12_ fraction, *r* and *k* are displayed on logarithmic colour scales. Release and uptake rates are geometric means across biological replicates in the absence of added glucose and were obtained from the first-order models. Trait-group ordinals correspond to those used in Fig. 4, 5 S12, S13, S14.

**Fig. S17.**
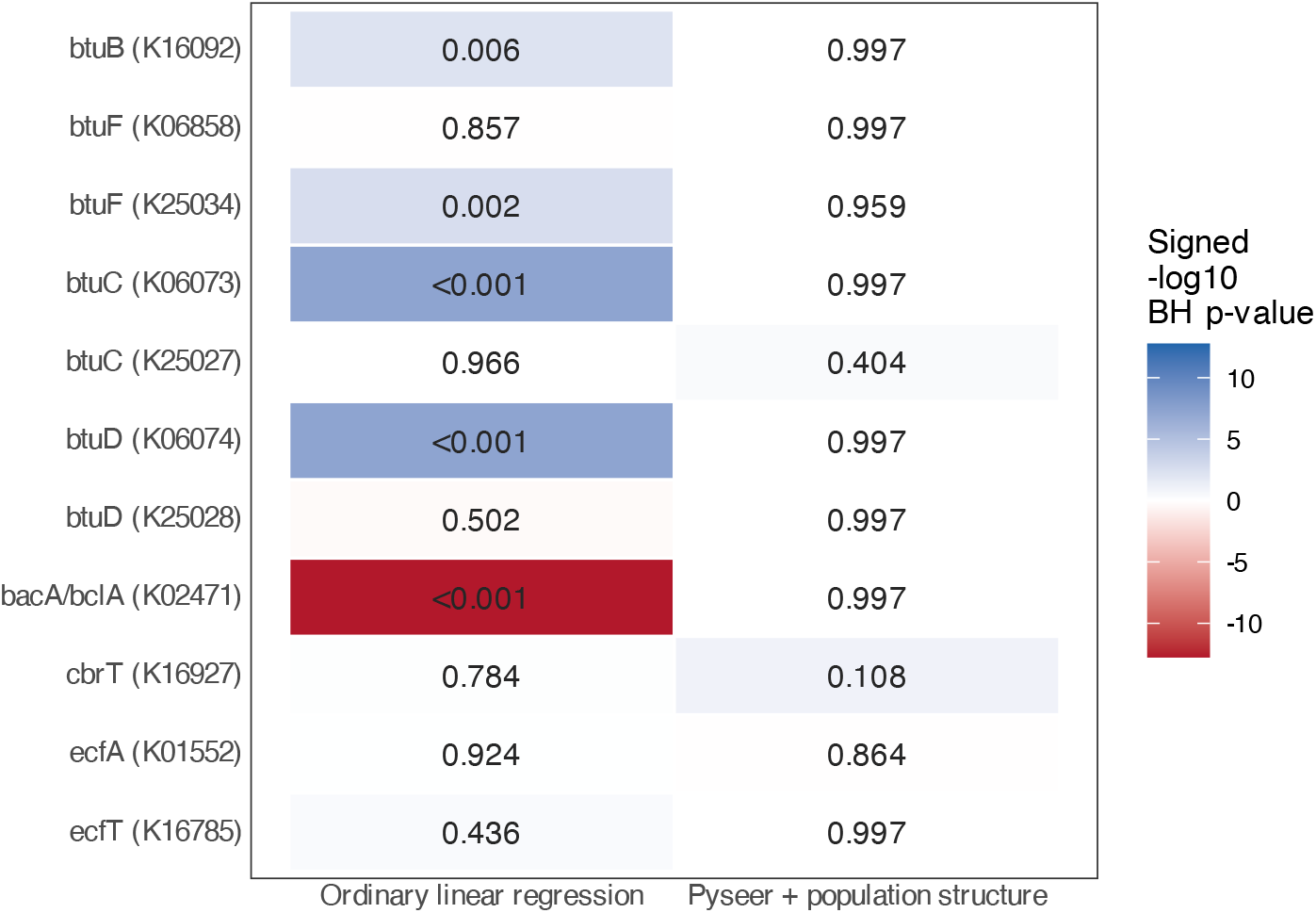
Associations between candidate B_12_-transporter KOs and strain-level mean B_12_ uptake. KO copy counts were binarized as absent (count = 0) or present (count *>* 0). For the ordinary linear-regression analysis, mean B_12_ uptake was modelled as a function of this binary indicator; the fitted coefficient therefore represents the difference in mean uptake between strains in which the KO was present and absent. These results were compared with likelihood-ratio tests from Pyseer linear mixed models, which tested the same presence–absence associations while accounting for population structure. Benjamini–Hochberg adjustment was performed across all variable genomic KOs tested by each method, rather than only across the 11 candidate transporters displayed here. Tile colour represents the signed − log_10_ Benjamini–Hochberg-adjusted *P* -value, with the sign determined by the fitted association coefficient; values printed within tiles are the adjusted *P* -values. Some KOs had perfectly correlated presence–absence profiles across strains (btuC-K06073 and btuD-K06074). Pyseer tested one representative from each such cluster, and the representative’s coefficient and *P* -value were assigned to the other cluster members for display; correlated members were not counted as additional independent tests during multiple-testing correction.

**Fig. S18.**
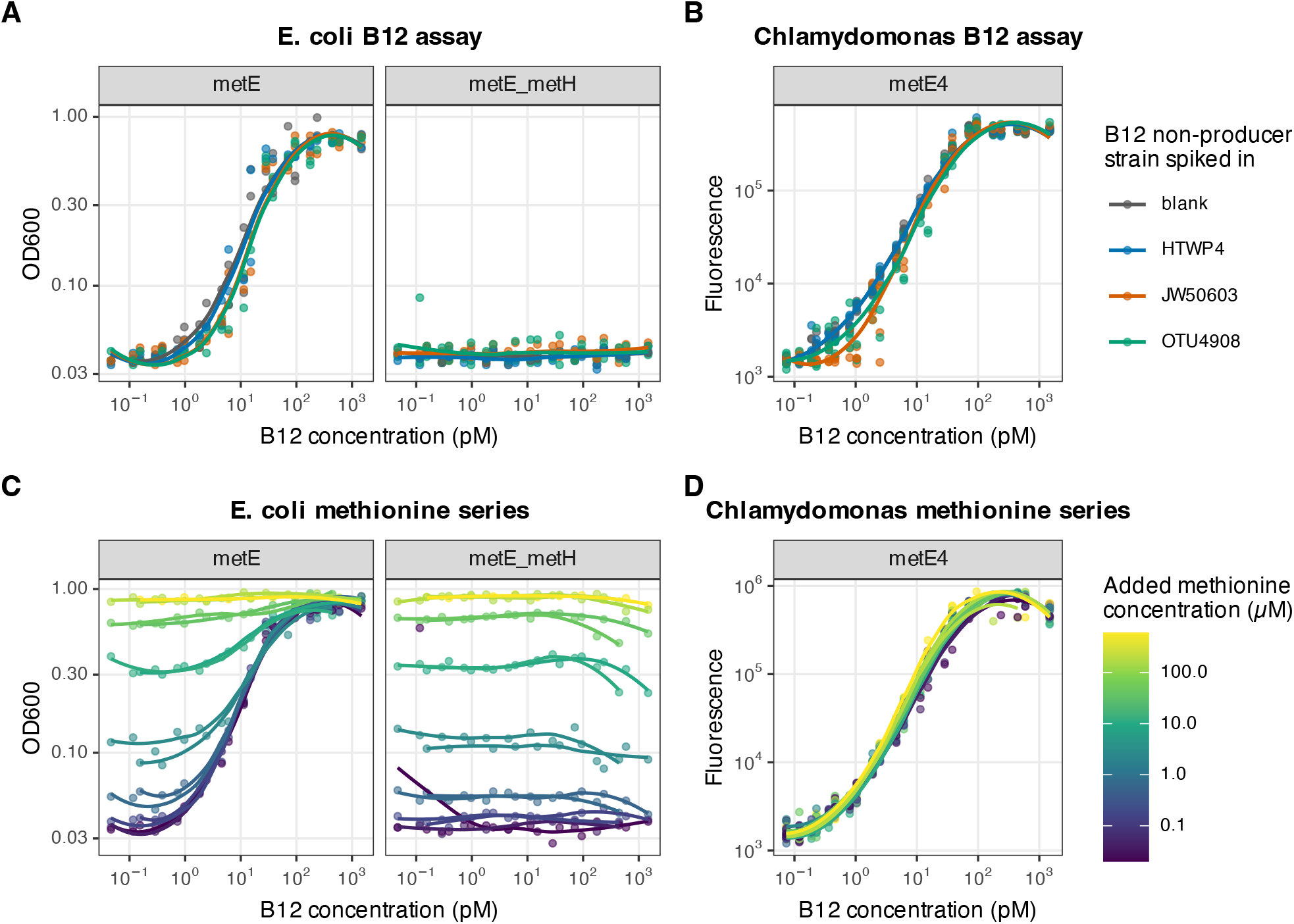
Comparison of the responses of *Escherichia coli* - and *Chlamydomonas*-based B_12_ bioassays to B_12_, methionine and bacterial sample matrices. Growth of the *E. coli* assay strains was quantified as optical density at 600 nm, whereas growth of the *Chlamydomonas metE4* assay strain was quantified using chlorophyll-pigment fluorescence, with excitation centred at 400 nm with a 100-nm bandwidth and emission centred at 700 nm with a 100-nm bandwidth. (A) Responses of the *E. coli metE* and *metE metH* strains to B_12_ standards prepared with no added bacteria or with the indicated non-B_12_-producing bacterial strain. (B) Response of *Chlamydomonas metE4* to the same B_12_ concentration series and bacterial treatments. (C,D) Responses of the *E. coli* and *Chlamydomonas* assay strains, respectively, to combinations of B_12_ and methionine in the absence of added bacteria; colour denotes the added methionine concentration. The *E. coli metE* strain can be supported by either B_12_, through the remaining B_12_-dependent methionine synthase MetH, or by exogenous methionine, whereas the *metE metH* double mutant responds only to methionine. In contrast, *Chlamydomonas metE4* responds to B_12_ but not to exogenous methionine. Points show individual measurements and lines show locally fitted response curves.

**Fig. S19.**
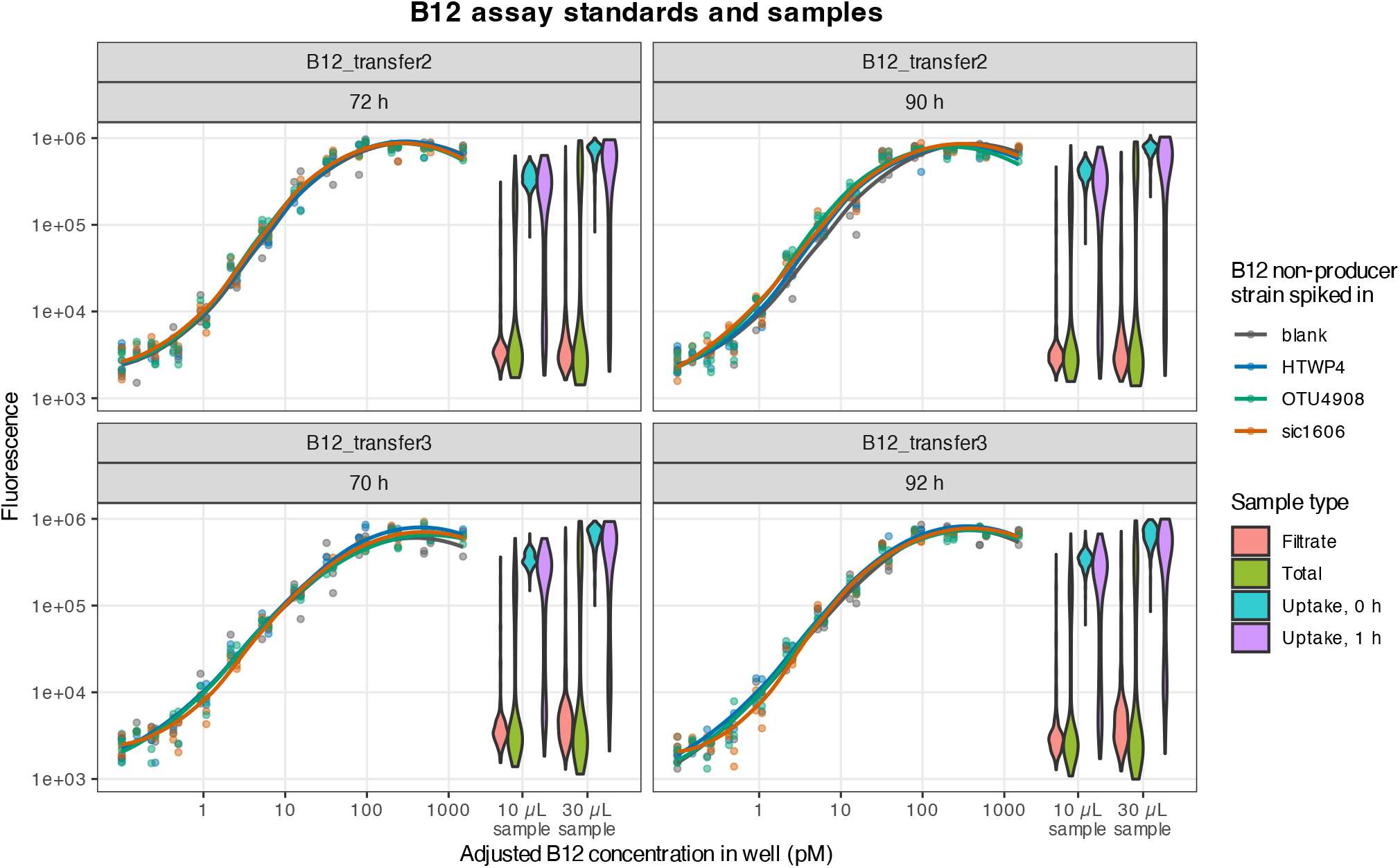
B_12_-bioassay standard curves and sample fluorescence distributions for the original B_12_ trait survey of 274 unique bacterial strains, from which the phenotypes analysed in Figures 1–3 were derived. Coloured points and fitted curves show B_12_ standards prepared in blank medium or in culture filtrate from the indicated non-B_12_-producing bacterial strain, allowing potential sample-matrix effects to be assessed. Violin plots show the fluorescence distributions of filtrate, total-culture, uptake-baseline and uptake-after-1-h samples. Each sample was assayed at two input volumes, 10 and 30 *µ*L, in a final assay volume of 100 *µ*L. The top and bottom facet rows represent two independent experimental replicates. The left and right columns show the two measurement times for each B_12_ assay: 72 and 90 h for the first replicate and 70 and 92 h for the second replicate, respectively. Both axes are shown on logarithmic scales.

## 6 Supplementary tables

**Table S1**. Bacterial strain metadata and GTDB-Tk taxonomy.

**Table S2**. Media and stock-solution recipes.

**Table S3**. Selected experimental strains, GTDB-Tk taxonomy, B_12_ trait groups, and strain labels used in Figures 4 and 5 and the associated supplementary figures.

## Acknowledgements

We thank members of the Kuehn laboratory for guidance and discussion on the project throughout its development. We also thank Alison Smith and Lorraine Archer for providing the B_12_-dependent Chlamydomonas reinhardtii strain used here for the B_12_ bioassays, as well as Sam Light and Josh Stemczynski for the *Escherichia coli* K-12 MG1655-derived B_12_ bioassay strains, which were originally provided by the USDA-ARS Culture Collection (NRRL).

## Author contributions

F.B. and S.K. conceived the study. F.B. designed the experimental work, performed the experiments, and conducted the primary data analyses. T.J. performed genome annotation and comparative genomic analyses and constructed the phylogenetic trees. P.M. performed 16S rRNA gene sequencing of the isolates and the B_12_-supplementation growth experiments. K.S. performed most of the marine bacterial strain isolations and associated media preparation and curated the resulting data. J.Z. performed the *pyseer* analyses. S.G., F.B., and S.K. developed the B_12_-flux models and generated the associated predictions. C.P. provided marine bacterial strains. C.D. supervised and contributed to the genotype–phenotype association and prediction analyses. M.M., C.P., and C.D. contributed to the conceptual development of the study. S.K. supervised the overall study. F.B. wrote the original manuscript. F.B. and S.K. revised the manuscript with input from all authors. All authors reviewed and approved the final manuscript.

## Declarations

### Generative AI disclosure

OpenAI Codex was used to assist with code review and modification, reproducibility checks, repository documentation, and editorial review of manuscript materials. All AI-assisted outputs were reviewed and verified by the authors, who take responsibility for the content of the manuscript and the associated analyses.

